# FAM111A regulates replication origin activation and cell fitness

**DOI:** 10.1101/2020.04.22.055574

**Authors:** Diana O. Rios-Szwed, Elisa Garcia-Wilson, Luis Sanchez-Pulido, Vanesa Alvarez, Hao Jiang, Susanne Bandau, Angus Lamond, Chris P. Ponting, Constance Alabert

## Abstract

FAM111A is a replisome associated protein and dominant mutations within its trypsin-like peptidase domain are linked to severe human developmental syndromes. However, FAM111A functions and its putative substrates remain largely unknown. Here, we showed that FAM111A promotes origin activation and interacts with the putative peptidase FAM111B, and we identified the first potential FAM111A substrate, the suicide enzyme HMCES. Moreover, unrestrained expression of FAM111A wild-type and patient mutants impaired DNA replication and caused cell death only when the peptidase domain remained intact. Altogether our data reveal how FAM111A promotes DNA replication in normal conditions and becomes harmful in a disease context.

## INTRODUCTION

FAM111A is expressed in all human tissues and has been initially proposed to play a role in tumorigenesis and viral host range restriction (Akamatsu et al., 2012; Fine et al., 2012). In humans, heterozygous point mutations in FAM111A are linked to two severe developmental syndromes: the Kenny-Caffey syndrome (KCS2, OMIM-127000) and Gracile Bone Dysplasia (GCLEB, OMIM-602361) characterized by, among others, short stature, hypoparathyroidism and dense or gracile bones. Remarkably, the R569H point mutation in the FAM111A gene is found in seven unrelated KCS2 patients, supporting a causal effect of FAM111A mutation in in KCS2 (Abraham et al., 2017; Isojima et al., 2014; Unger et al., 2013) (Fig. 1A). Although FAM111A catalytic activity has not been shown *in vitro*, recent work revealed that *in vivo* FAM111A exhibits autocleavage activity when its peptidase domain is intact (Kojima et al., 2020), strongly supporting that FAM111A is a peptidase. Interestingly, the R569H mutation, and those of three other KCS2 and GCLEB patients, Y511H, S342Del and D528G, do not compromise but rather enhance FAM111A autocleavage activity (Kojima et al., 2020). As FAM111A function and substrates remain unknown, it is unclear how gain-of-function mutations contribute to KCS2 and GCLEB etiology. To provide better diagnosis and management of these conditions, it is therefore fundamental to understand the function of FAM111A in normal and disease contexts.

**Figure 1.**
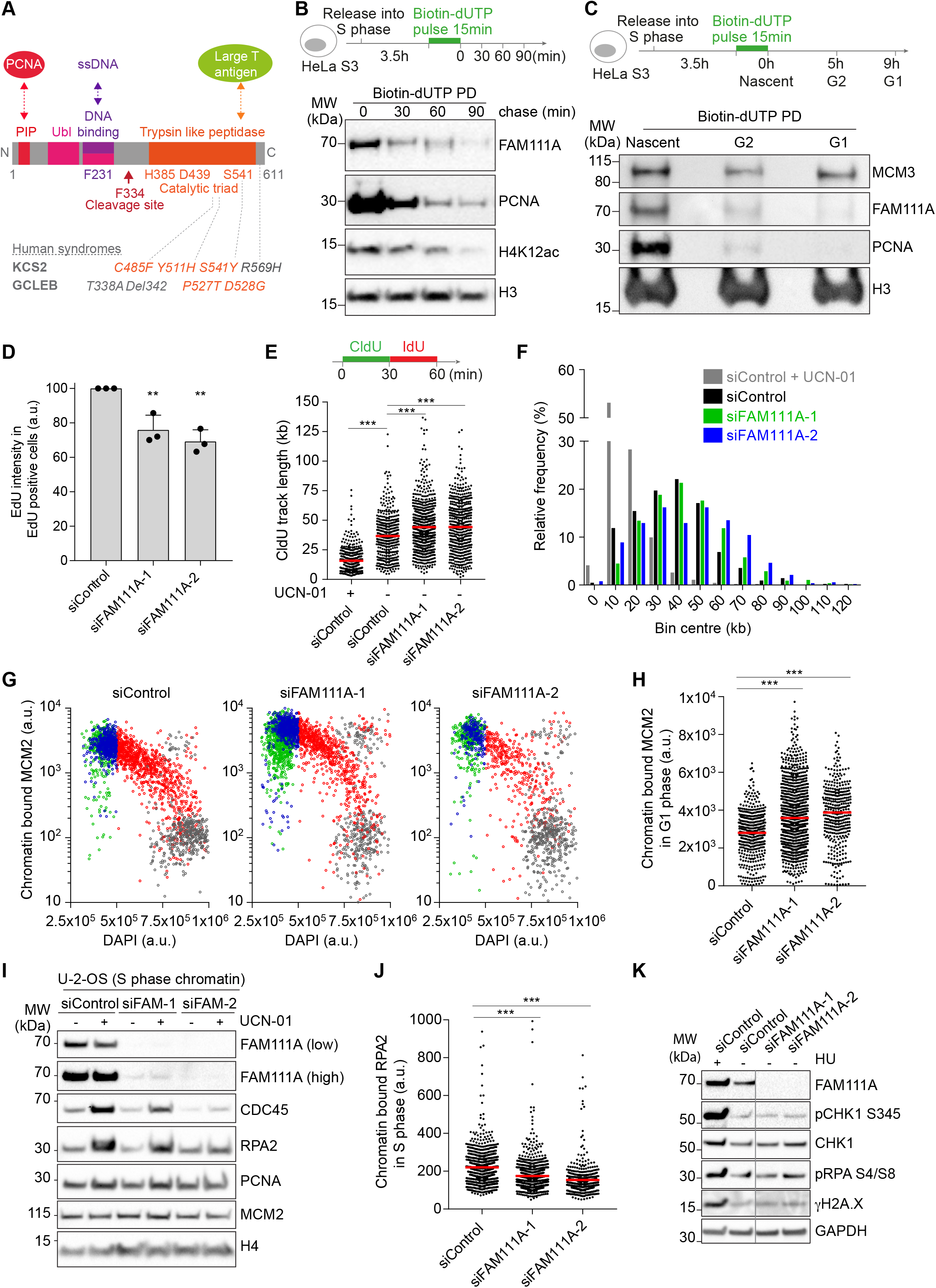
FAM111A promotes origin activation at the G1/S transition. **A**. Schematic representation of FAM111A domain structure with notable residues and direct interactors highlighted. **B, C**. NCC analysis of FAM111A recruitment to replicated chromatin in S phase (B), and the following G2 and G1 phase (C). **D**. DNA synthesis measured by EdU incorporation. U-2-OS cells were transfected with the indicated siRNAs and after 48 h pulse labelled with 40 μM EdU for 20 min. Data are represented as mean with standard deviation (S.D.) of three independent experiments; 785 < n < 2656 cells were analyzed per condition and per experiment. **E**. Analysis of replication fork speed by DNA combing. Top, labelling strategy. Bottom, size distribution of CldU track length. Red bar represents the median; n > 421 tracks were analyzed. **F**. Frequency distribution of CldU track length from E. **G**. Chromatin bound MCM2 levels in U-2-OS cells shown as a function of DAPI intensity and cell cycle stage detected by HTM. Cell cycle gating shown in Fig. S1E. Green, G1 phase; Blue, early S phase; Red, mid/late S phase; Grey, G2/M phase. From left, n = 2317, 2335, 1424. **H**. Quantification of chromatin bound MCM2 in G1 phase analyzed in G. From left, n = 474, 999,380. **I**. Immunoblot of chromatin fractions from cells synchronized in S phase and treated with 300 nM UCN-01 for 2 h. Results for asynchronous cells is shown in Fig. S1N **J**. Chromatin binding of RPA2 in S-phase cells detected by HTM. S phase cells were gated based on chromatin bound PCNA intensities as in Fig. S1O. From left, n = 910, 931, 453. **K**. Immunoblot of whole cell extracts from siRNA transfected U-2-OS cells. Data are representative of two (B, C, E, F), and three (D, G-K) independent experiments. siControl, non-targeting siRNA; a.u., arbitrary units. D, unpaired Student’s t-test. E, H, J, Mann-Whitney test. ***P < 0.001, ** P < 0.01.

Pioneering work suggested that FAM111A functions as a viral host range restriction factor (Fine et al., 2012) as upon SV40 viral infection, FAM111A is recruited to sites of viral replication and reduces viral replication rates (Fine et al., 2012; Tarnita et al., 2019). Similarly, FAM111A is recruited to cellular DNA replication sites and its transient overexpression blocks DNA replication (Alabert et al., 2014; Tarnita et al., 2019). However, in absence of FAM111A, the rate of DNA synthesis is also reduced suggesting that FAM111A may also play a positive role in DNA replication (Alabert et al., 2014). Consistent with this, FAM111A has recently been shown to promote fork progression through artificially created DNA binding protein crosslinks (DPC) (Kojima et al., 2020). Mechanistically, FAM111A binds directly to proliferating cell nuclear antigen (PCNA) through an N-terminal PCNA interacting protein box (PIP) (Alabert et al., 2014), and to single-stranded DNA (ssDNA) through a newly identified ssDNA binding domain (Kojima et al., 2020) (Fig. 1A). Thus, clues have emerged for possible new roles for FAM111A under stress conditions, yet the molecular function of FAM111A under normal conditions remains unclear. Moreover, the repertoire of FAM111A substrates has yet to be identified.

Here, we have investigated the molecular mechanisms that link FAM111A to DNA replication. We report that under normal and replication stress conditions, FAM111A promotes activation of licensed replication origins. FAM111A interacted with FAM111B, which together with FAM111A promoted DNA replication, while also having FAM111A independent role(s). Overexpression of FAM111A or expression of FAM111A harboring KCS2 and GCLB2 patient mutations caused DNA damage and cell death which were dependent of an intact FAM111A peptidase domain, and partially contingent on FAM111A recruitment to PCNA. Finally, our results identified the first two potential substrates of FAM111A, the suicide enzyme HMCES and the ribosomal protein RPL26L.

## RESULTS AND DISCUSSION

### FAM111A promotes origin activation at the G1/S transition

To uncover the role of FAM111A in DNA replication, we first examined FAM111A recruitment to newly replicated chromatin by nascent chromatin capture (NCC). Consistent with our previous work (Alabert et al., 2014), FAM111A was enriched on newly replicated chromatin (Fig. 1B, Fig. S1A). The majority of FAM111A dissociated from replicated chromatin within the first 30 minutes after DNA synthesis, mirroring the behavior of replisome components. To further understand FAM111A recruitment to chromatin, we chased replicated chromatin through the following G2 and G1 phases (Fig. 1C, S1B, C). Unlike MCM3 which was loaded *de novo* onto chromatin in G1 phase, FAM111A did not re-associate with chromatin for the rest of the cell cycle, suggesting that FAM111A’s principal function on chromatin is during DNA replication.

Next, we monitored the ability of cells to proliferate and replicate in absence of FAM111A (Fig. S1D). FAM111A depleted U-2-OS cells accumulated in G1 phase (Fig. S1E, F) and exhibited impaired cell proliferation (Fig. S1G, H). Furthermore, the rate of DNA synthesis was reduced in these cells (Fig. 1D), consistent with previous observations using independent siRNAs (Alabert et al., 2014). To determine whether the reduced DNA synthesis rate resulted from a replisome progression defect (slower forks) or a replication initiation defect (fewer forks), we analyzed DNA replication at the single molecule level using DNA molecular combing. To this end, newly replicated DNA was successively pulse labeled using two nucleotide analogs CldU and IdU, and CldU signals were used to determine replisome elongation rates. Replisome progression was not affected upon FAM111A depletion, with fork speed being slightly increased instead (Fig. 1E, F). In contrast, the inter-fork distance was slightly increased in FAM111A depleted cells (Fig. S1I), suggesting that under these conditions fewer origins had initiated. We further tested this hypothesis by artificially triggering dormant origin activation with the CHK1 inhibitor 7-hydroxystaurosporine (UCN-01), and measured the resulting inter fork distance (Feng et al., 2016; Ge et al., 2007; Maya-Mendoza et al., 2007; Petermann et al., 2010; Saldivar et al., 2017). As expected the inter fork distance was reduced in control cells upon UCN-01 treatment due to the activation of dormant origins (Fig. S1I, J). In FAM111A depleted cells however, the inter-fork distance remained higher than in control cells (Fig. S1K). Moreover, in absence of FAM111A the induction of origin firing upon UCN-01 treatment was also less efficient compared to control (Fig. S1L), suggesting that dormant origin firing is also impaired. Altogether, these data suggest that FAM111A promotes DNA replication initiation at both active and dormant origins, while being dispensable for fork progression.

DNA replication initiation is a two-step process. In G1 phase, origins are licensed by the loading of MCM2-7 complexes, while in S phase, a fraction of the origins are activated by the CDK and DDK dependent recruitment of CDC45, the GINS complex, and the rest of replisome (Ganier et al., 2019; Marchal et al., 2019). To identify at which stage of replication initiation FAM111A may function, we first examined the origin licensing efficiency in FAM111A depleted cells by quantifying MCM2 abundance on chromatin in G1 phase cells by high throughput microscopy (HTM) (Fig. 1G, S1E). In FAM111A depleted cells, MCM2 loading was not impaired (Fig. 1G, H), indicating that FAM111A does not promote origin licensing. In contrast, CDC45 abundance on chromatin was reduced upon FAM111A depletion (Fig. 1I, S1M, N), suggesting that FAM111A may promote origin firing. Mirroring CDC45, chromatin bound RPA levels were also reduced in S phase upon FAM111A depletion (Fig. 1J, S1N, O) while the pool of nuclear RPA was unaffected (Fig. S1P). Moreover, consistent with the ability of FAM111A to promote dormant origin activation, CDC45 recruitment to chromatin was also impaired in UCN-01 treated FAM111A deficient cells (Fig. 1I, S1N). Importantly, FAM111A depletion did not activate the ATR-CHK1 pathway (Fig. 1K), excluding that in FAM111A depleted cells, origin activation was impaired indirectly through activation of the ATR-CHK1 pathway (Saldivar et al., 2017). Altogether, these data indicate that FAM111A promotes activation of licensed origins. Mechanistically, as FAM111A is expected to possess a peptidase activity, it suggests that FAM111A may degrade factors that inhibit origin firing, either directly as the RIF1-PP1 complex (Dave et al., 2014), or indirectly by targeting chromatin based processes controlling access of firing factors to DNA (Feng et al., 2016; Wu et al., 2017).

### FAM111A depletion protects cells against replication stress

As firing of dormant origins is essential during replicative stress to ensure completion of DNA replication (Saldivar et al., 2017), we next determined whether FAM111A may play a role during replication stress. We used two DNA replication inhibitors, hydroxyurea (HU) which blocks the ribonucleotide reductase RNR and therefore deoxynucleotide production, and the DNA polymerase-α inhibitor aphidicolin (APH). Both drugs rapidly impair replisome progression while allowing dormant origin firing, collectively leading to accumulation of ssDNA and recruitment of RPA (Fig. S2A). Here, we assessed the drug induced accumulation of RPA onto chromatin upon depletion of FAM111A using HTM. As expected, RPA accumulation on chromatin was detectable 2 hours after HU treatment (Fig. 2A, B). However, chromatin bound RPA levels were markedly reduced in FAM111A deficient cells compared to control (Fig. 2A, B). Similar results were observed upon APH treatment (Fig. 2C), in HU treated cells transfected with distinct set of siRNAs (Fig. S2B, C) and in cells stably expressing GFP-RPA1 (Fig. S2D). These data further support that FAM111A promotes dormant origin activation, and suggest that FAM111A may play an additional role in ssDNA exposure at stalled forks.

**Figure 2.**
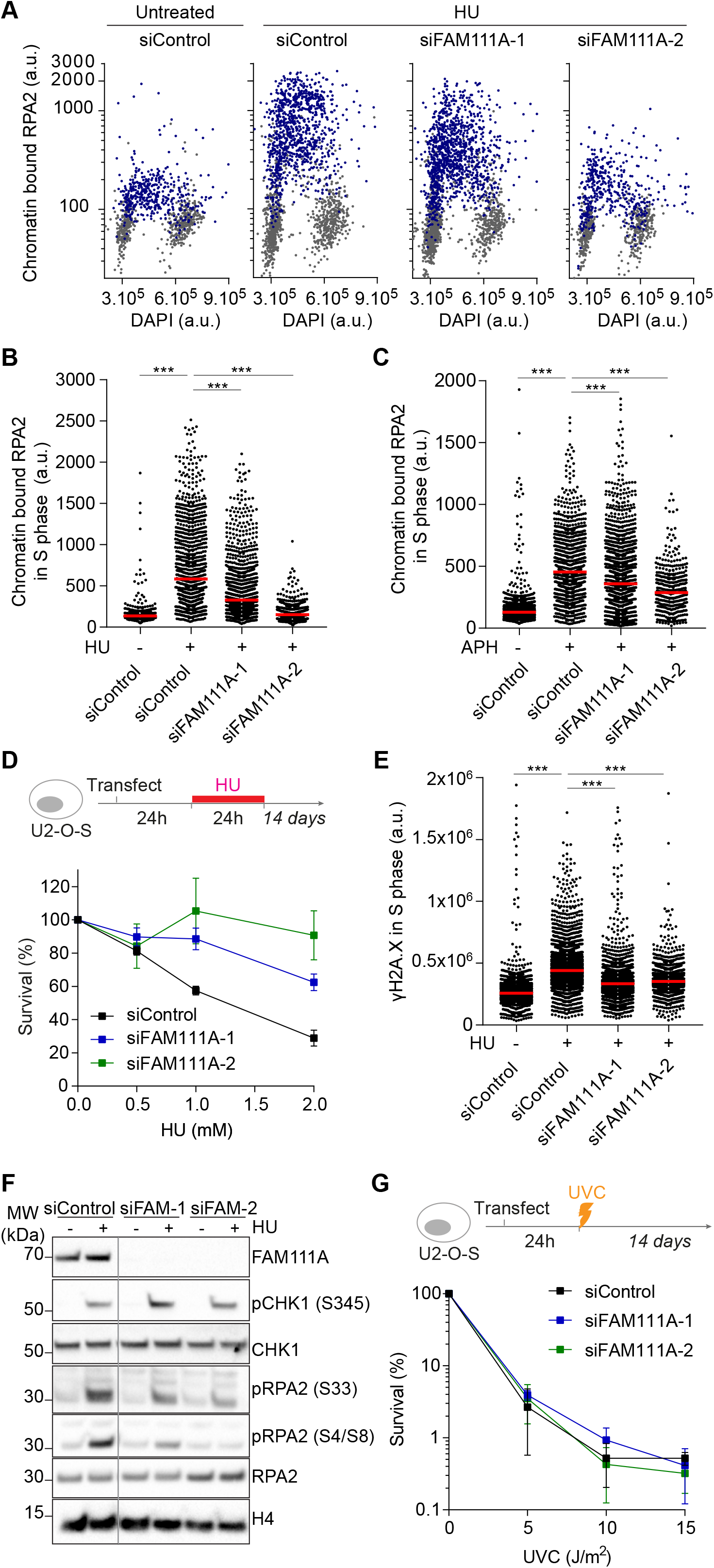
FAM111A depletion protects cells against replicative stress. **A**. Chromatin bound RPA2 intensity in U-2-OS cells treated with 3 mM HU for 2 h and detected by HTM. Data shown as a function of DAPI intensity. Gating strategy as in Fig. S1O. Blue, PCNA positive; Grey, PCNA negative. **B**. Quantification of chromatin bound RPA2 in S phase cells analyzed in A. From left, n = 457, 991, 1216, 457. **C**. Chromatin binding of RPA2 in S phase cells treated with 50 μg/mL APH for 2 h, analyzed as in A. From left, n= 1494, 999, 1349, 402. **D**. Clonogenic survival assays of siRNA-transfected cells treated with HU as indicated. **E**. Chromatin abundance of γH2A.X in S phase analyzed as in A. From left, n= 2720, 2486, 1664, 1149. **F**. Immunoblot of whole cell extracts from siRNA-transfected cells treated as in A. **G**. Clonogenic survival assays of siRNA-transfected cells treated with UVC as indicated. Data are representative of three (A-C, F) and two (D, E, G) independent experiments. B, C, E, Mann-Whitney test, ***P < 0.001.

We next examined the role of FAM111A during prolonged replication stress by exposing siRNA transfected cells to a 24-hour HU treatment and monitoring their survival. Interestingly, FAM111A depleted cells were resistant to HU (Fig. 2D) suggesting that FAM111A depletion protects cells against HU induced replication stress. HU resistance has previously been observed in cells with lower respiration rate (Nakayashiki and Mori, 2013) or cells with a lower number of active forks exposed to replicative stress (Feng et al., 2016). As FAM111A promotes origin firing, it is possible that in FAM111A depleted cells fewer forks were exposed to HU. Supporting this, lower levels of DNA damage were observed in absence of FAM111A after acute HU treatment (Fig. 2E) although the ATR-CHK1 pathway remained functional in these cells (Fig. 2F, S2E).

Finally, as FAM111A binds directly to PCNA, we tested whether FAM111A may play a role in nucleotide excision repair (NER), a PCNA dependent repair pathway. FAM111A depleted cells were not more sensitive to short wavelength UV (UVC) than control cells (Fig. 2G) suggesting that FAM111A may not be required for NER repair. Interestingly, FAM111A is strongly ubiquitylated upon UVC treatment (Povlsen et al., 2012). Ubiquitination may thus provide a mechanism to trigger FAM111A eviction from sites of repair where PCNA is involved.

### Unrestrained FAM111A peptidase activity at replication fork interferes with DNA replication

In cancer, no mutation hot spot has yet been identified in the FAM1111A gene. Instead, FAM111A is often overexpressed (COSMIC GRCh38.v90). Thus, we dissected the impact of FAM111A overexpression on DNA replication and genome stability. Stable cell lines conditionally expressing FLAG-HA-FAM111A wild type (WT) and FLAG-HA-FAM111A mutants were generated (Fig. 3A). FAM111A WT overexpression increased γH2A.X levels (Fig. 3B, S3A-C) and cell death (Fig. 3C), while reducing DNA synthesis rates (Fig. 3D). Single cell analysis revealed that 40% of cells in mid-S phase had no EdU incorporation (Fig. 3E, S3E), suggesting that in these cells, replication forks may have stopped and collapsed. Moreover, we observed a clear anti-correlation between chromatin bound FAM111A levels and EdU incorporation (Fig. 3F). Consistent with this, EdU levels could be partially rescued by depleting endogenous FAM111A (Fig. 3G), further supporting a direct link between DNA synthesis and FAM111A abundance. Interestingly, FAM111A accumulated onto chromatin and blocked cells preferentially at the G1/S transition (Fig. 3H, S3D). Altogether these data indicate that FAM111A overexpression impairs DNA synthesis, induces DNA damage and is deleterious to cell fitness.

**Figure 3.**
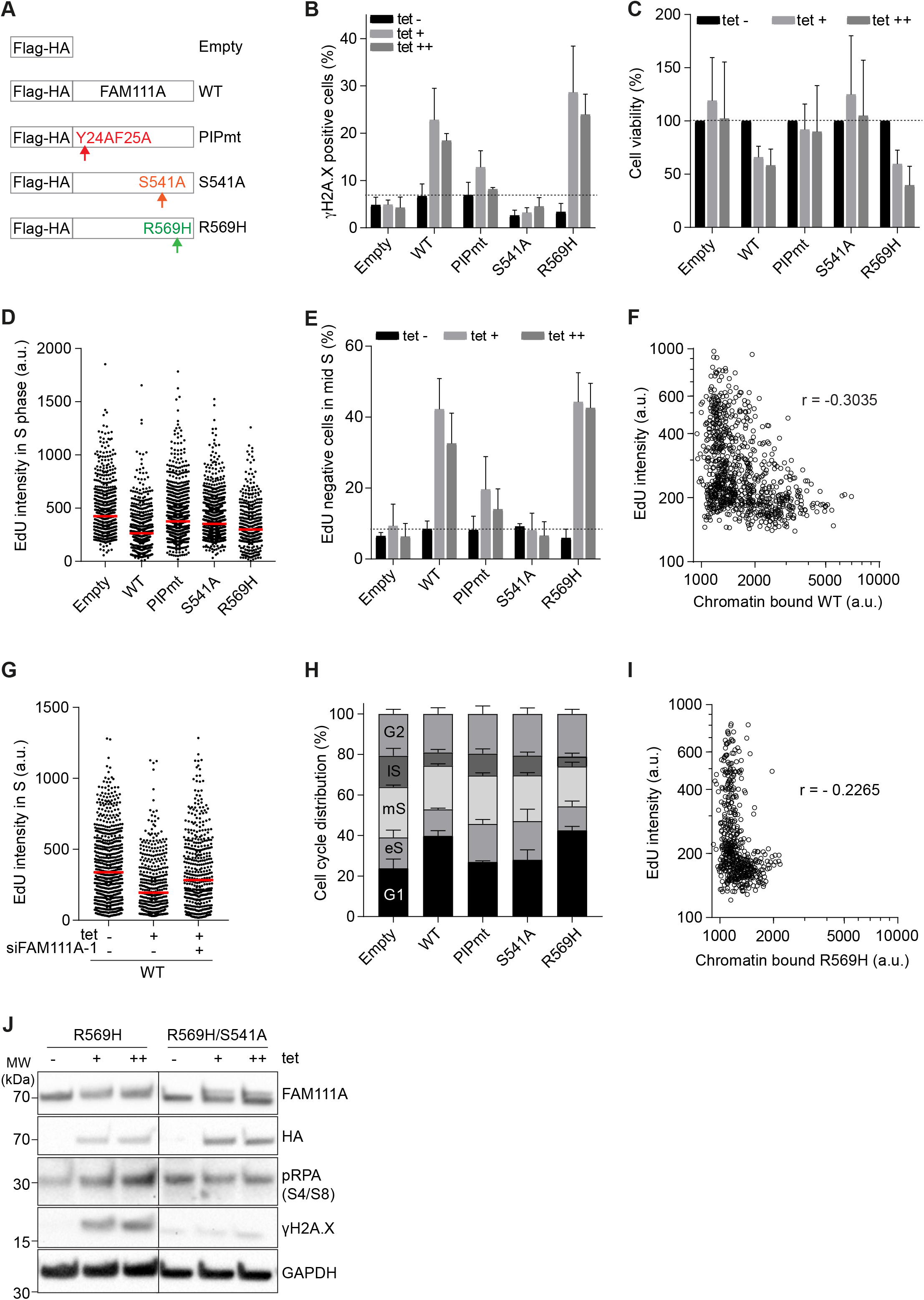
Unrestrained FAM111A peptidase activity is toxic for cell fitness. **A**. Schematic representation of tetracycline (tet) inducible Flag-HA-FAM111A constructs. **B**. Quantification of γH2AX positive cells upon Flag-HA-FAM111A overexpression. Gating strategy shown in Fig. S3B, C. **C**. Quantification of cell numbers 24 h after Flag-HA-FAM111A overexpression relative to uninduced condition. **D**. EdU intensities in S phase upon Flag-HA-FAM111A overexpression induced with 0.5 μg/mL tet. Cells were gated as in Fig. S3C. From left, n = 545, 529, 787, 973, 392. **E**. Quantification of EdU negative cells in mid S phase upon Flag-HA-FAM111A overexpression. Cells were gated as in Fig. S3E. **F**. EdU intensities shown as a function of chromatin bound FH-FAM111A in pre-extracted U-2-OS cells. r, Pearson correlation. n = 918. **G**. EdU intensities of siRNA-transfected cells and treated with 1 μg/mL tet for 24 h, as indicated. From left, n = 653, 417, 444. **H**. Cell cycle distribution upon Flag-HA-FAM111A overexpression. Cells were gated as in Fig. S3C. **I**. EdU intensities shown as a function of chromatin bound Flag-HA-FAM111A-R569H in pre-extracted U-2-OS cells. r, Pearson correlation. n = 584. **J**. Immunoblot of whole cell extracts from asynchronous cells 24 h after tetracycline induction. For B, C, E, and H, data are represented as mean + SD of n = 3 experiments. Data are representative of three (D, F, I, J) and two (G) independent experiments. tet (−), uninduced cells; tet (+), 0.5μg/mL tetracycline; tet (++), 1.0 μg/mL tetracycline.

FAM111A is predicted to be a serine trypsin-like peptidase, harboring in the C-terminus a conserved catalytic triad of histidine (385), aspartate (439), and serine (541) (Fig. 1A, S3F). We generated a FAM111A predicted peptidase dead mutant inducible cell line by replacing serine 541 by an alanine (S541A) (Fig. 3A). In this mutant, the serine-OH group that acts as the nucleophile attacking the peptide bond of the substrate is missing. Unlike FAM111A-WT, expression of S541A did not increase level of γH2AX (Fig. 3B) and did not cause cell death (Fig. 3C) or cell cycle arrest (Fig. 3H). These results revealed that the increased abundance of the FAM111A peptidase activity specifically is detrimental to cells. Notably, expression of FAM111A PIP mutant (PIPmt) induced lower levels of γH2A.X and cell death compared to FAM111A-WT (Fig. 3B, C), and showed higher DNA synthesis rates (Fig. 3D). As FAM111A binds to PCNA *in vitro* (Alabert et al., 2014), one possibility is that the cytotoxicity observed upon unrestrained peptidase activity is in part due to cleavage of one or more replisome components. Collectively, these results indicate that FAM111A abundance must be tightly controlled as both depletion and overexpression impair DNA synthesis and cell cycle progression. Moreover, unlike FAM111A depletion, FAM111A overexpression is cytotoxic and was dependent on an intact putative catalytic triad suggesting that excessive degradation of FAM111A substrates is deleterious for cell fitness.

### FAM111A function in disease etiology

To understand the molecular basis of KCS2’s etiology, we focused on R569H, the most frequent mutation identified in patients (Fig. 1A). R569H is a dominant monoallelic mutation that appears to confer hyperactive peptidase activity (Kojima et al., 2020). To mimic KCS2 patients’ cells, we induced expression of Flag-HA-FAM111A-R569H in cells retaining endogenous FAM111A (Fig. 3A). FAM111A-R569H expression increased γH2A.X levels and cell death (Fig. 3B, C), and as for FAM111A-WT, DNA synthesis rates were reduced (Fig. 3D). Similarly, cells were blocked in mid-S phase with no EdU incorporation (Fig. 3E), suggesting that replication forks collapsed, and the correlation between chromatin bound R569H levels and EdU incorporation was even lower than for FAM111A-WT (Fig. 3I). Strikingly, in the double mutant S541A-R569H, where the peptidase activity is inactivated, γH2A.X levels were rescued (Fig. 3J), further supporting that unrestrained FAM111A peptidase activity is deleterious for cell survival.

FAM111A-R569H abundance was low compared to other generated FAM111A mutants (Fig. S3A, D, G) consistent with its enhanced autocleavage activity (Kojima et al., 2020). Similarly, despite its low abundance, R569H impact on DNA replication and cell viability was comparable to that of highly expressed WT-FAM111A, supporting that R569H is a gain-of-function mutation, producing a hyperactive form of the peptidase. Another KCS2 patient mutant, Y511H, which also exhibits enhanced autocleavage activity (Kojima et al., 2020) was also poorly expressed, as well as a previously unstudied GCLEB patient mutant, T338A (Fig. S3G). What is unique about this last mutation, and how it may lead to a distinct syndrome remain to be explored. A mutation of the catalytic triad’s serine (S541Y) has recently been found in a young KCS patient (Abraham et al., 2017). Although, it is unknown how this mutation affects FAM111A catalytic activity, all disease mutations tested so far have performed as gain-of-function mutations, indicating that the level of FAM111A peptidase activity must be tightly regulated to allow proper cell proliferation and organismal development.

Notably, although FAM111A disease mutants exhibit enhanced autocleavage ability, KCS2 patients’ cells may not be primarily affected by the drop in FAM111A abundance as FAM111A depletion did not lead to increased DNA damage or substantial cell death (Fig. 1K). Instead, unrestrained FAM111A peptidase activity may lead to hyper degradation of FAM111A substrates or degradation of additional substrates. The latter may explain how FAM111A can play a positive role in DNA replication, promoting origin firing, and becomes harmful in a disease context.

### Identification of FAM111A binding partners and putative substrates

FAM111A autocleavage site suggests a chymotrypsin like peptidase specificity (Kojima et al., 2020). Predicting its substrates *in silico* is unlikely as protease substrate specificities are often broad and highly dependent on amino acid sequence and tertiary structure (Goettig et al., 2019). Therefore, to identify potential FAM111A substrates we combined two strategies. First, we performed an in depth FAM111A interactome analysis using affinity purification and mass spectrometry (AP-MS) of endogenous FAM111A from whole cell extracts and chromatin fractions. Chromatin was isolated using a no salt buffer as described in (Mendez and Stillman, 2000) to preserve charge-based interaction with DNA or histones (Fig. S4A-C). Buffer containing nonionic detergent and salt (Saredi et al., 2016) disrupted FAM111A binding to chromatin, suggesting that the majority of FAM111A binds loosely to chromatin and / or predominately to highly accessible chromatin (Henikoff et al., 2009), consistent with its binding to replicating chromatin (Fig. S4A, B). In whole cell and chromatin extracts, FAM111A’s top interactor was FAM111B (Fig. 4A, B, Table S1). The most enriched replisome component was RFC1 in whole cell extracts and MCM6 on chromatin fractions although these enrichments were non-significant.

**Figure 4.**
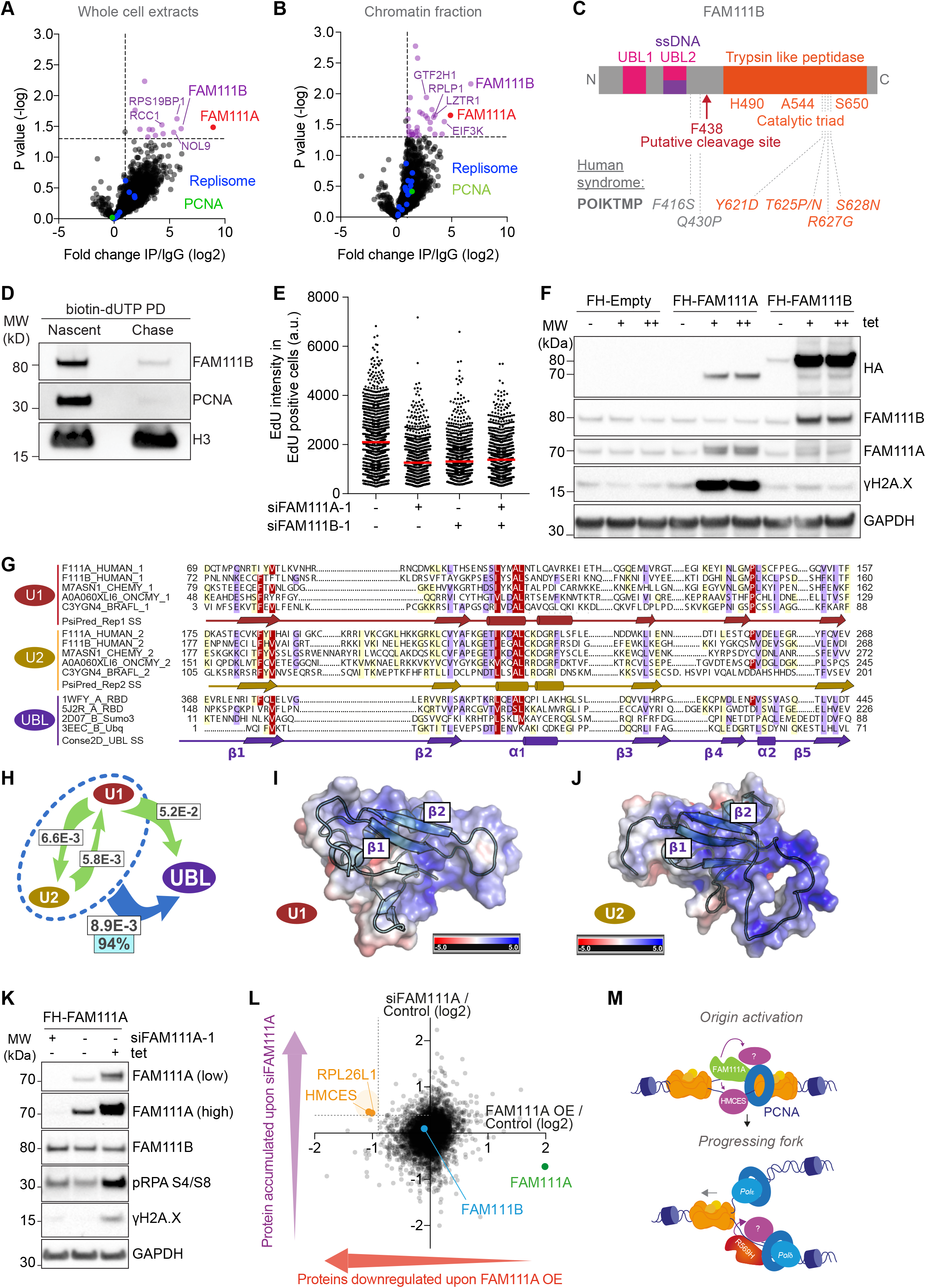
Identification of FAM111A binding partners and putative substrates. **A, B.** FAM111A complexes from whole cell extract (A) and chromatin fraction (B). **C**. Schematic representation of FAM111B domain structure and notable residues. **D**. NCC analysis of FAM111B recruitment to nascent chromatin in HeLa S3 cells. Input showed in Fig. S4D. **E.** EdU intensity in EdU positive cells in siRNA transfected U-2-OS cells. n = 1307, 974, 909, 986. **F.** Immunoblot of whole cells extracts after induction of Flag-HA-FAM111A or Flag-HA-FAM111B. FH-Flag-HA; (+), 0.5 μg/mL tet; (++), 1 μg/mL tet. **G**. Multiple sequence alignment of two consecutive UBL domains in FAM111. Red, FAM111 UBL repeats 1 (U1); Yellow, FAM111 UBL repeats 2 (U2); Purple, selection of UBL domains with known structure (UBL). Secondary structure predictions were performed independently for U1 (PsiPred_Rep1 lane) and U2 (PsiPred_Rep2 lanes), and are consistent with UBL (Conse2D_UBL lane). α-helices, cylinders; β-strands arrows. Average BLOSUM62 score: red, > 1.5; violet, between 1.5 and 0.5; light yellow, between 0.5 and 0.2. **H.** HHpred analysis. White rectangles, HHpred profile-versus-profile comparison E-values from global profile search results. Arrows, profile search direction e.g., U1 aligns to U2 with E-value = 6.6×10_−3_. Dotted blue oval, HHpred searches against the PDB70 profile database using alignment of U1 and U2 repeats as input detected the UBL Ras-binding domain of mouse RGS14 (PDB ID: 1WFY) (UBL) with E-value of 8.9×10_−3_; Cyan rectangle, true-positive homology probability of 94%. **I J.** 3D models of FAM111A U1 and U2 repeats. Red, negative charge surface electrostatic potential; blue positive. **K**. Immunoblot of whole cell extracts from Flag-HA FAM111A inducible U-2-OS cells 48 h after siRNA transfection and 24 h after 1.0 μg/mL tetracycline induction. **L**. Ratio of FAM111A overexpression to control (x axis), and of siFAM111A to control (y axis) of proteins abundance determined by TMT mass spectrometry. **M.** Model of FAM111A function. FAM111A plays a positive role in DNA replication, promoting origin firing, and becomes harmful in a disease context. Upon origin activation, FAM111A degrades HMCES and additional factors to promote origin firing. In a disease context, the FAM111A-R569H patient grain-of-function mutant hyper-degrades FAM111A substrates or target additional factors, causing fork collapse and cell death. Data are representative of three (A), four (B) and two (D-F, K, L) independent experiments.

FAM111B is the paralog of FAM111A and also contains a trypsin like peptidase domain in the C-terminus (Fig. 4C). Mutations in FAM111B gene are associated with an autosomal dominant form of hereditary fibrosing poikiloderma (POIKTMP, OMIM-615704) (Fig. 4C). Interestingly FAM111B is produced in S phase and therefore has been suggested to play a role in DNA replication (Aviner et al., 2015). Like FAM111A, FAM111B localized transiently to newly replicated chromatin (Fig. 4D). FAM111B depletion also reduced DNA synthesis and, interestingly depletion of both FAM111A and FAM111B did not result in further EdU reduction (Fig. 4E). Furthermore, depletion of FAM111B and of both paralogs led to decreased RPA loading upon HU treatment (Fig. S4E). These results indicated that FAM111A and FAM111B may act in the same pathway to regulate DNA replication and that they do not compensate for one another. Moreover, depletion of FAM111A did not affect FAM111B protein and vice versa (Fig. S4F), indicating that although these proteins may interact, they may not cleave one another. Finally, unlike FAM111A, FAM111B overexpression did not increase γH2A.X levels (Fig. 4F), suggesting that FAM111B may have FAM111A independent roles. Supporting the latter, mutations in FAM111A and FAM111B lead to distinct human syndromes.

The ssDNA binding domain recently identified in FAM111A (Kojima et al., 2020) is well conserved in FAM111B (Fig. 4G) suggesting that FAM111B may also bind ssDNA. Interestingly, we identified in both paralogs two Ubiquitin-like (UBL) repeat domains, U1 and U2, which differ from each other mainly by the presence of a long positively charged loop rich in arginine and lysine between β-strands 1 and 2 in U2 (Fig. 4G-I). Notably, the ssDNA binding domain maps to the U2 (Fig. S3F). Given the vast ubiquity and functional diversity of the UBL fold (Kiel and Serrano, 2006) it is possible to imagine an evolutionary scenario in which these types of domains have been recruited to interact with ssDNA. Other UBL domains are known to interact with nucleic acids: SUMO-1 binds double stranded DNA (Eilebrecht et al., 2010) and SF3A1 UBL domain binds double stranded RNA (Martelly et al., 2019). To our knowledge, FAM111 UBL domain is the first case of a putative UBL domain that interacts with ssDNA.

As a second strategy to identify FAM111A putative substrates, we designed an unbiased approach based on TMT quantitative mass spectrometry. We analyzed the relative composition of whole cell extracts combining three conditions: untreated, siFAM111A and FAM111A overexpression (Fig. 4K). In this set-up, FAM111A substrate abundance is expected to be reduced upon FAM111A overexpression and enriched in siFAM111A condition. 6928 factors were identified and as expected, FAM111A was enriched in overexpressed conditions and reduced upon siRNA-based knockdown (Fig. 4L, Table S2). Factors upregulated in both FAM111A overexpression and siFAM111A compared to control were involved in DNA replication and apoptosis, supporting the observed cell accumulation in G1/S and its cytotoxicity (Fig. S4G). FAM111B abundance was unaffected (Fig. 4L), consistent with our previous observations (Fig. S4F), and further supporting that FAM111B may be a cofactor of FAM111A rather than a substrate. Replisome components were not significantly affected by FAM111A deregulation (Fig. S4H), suggesting that FAM111A may not target core replisome components, but replication accessory factors instead.

Only two factors qualified as FAM111A substrates: HMCES and RPL26L (Fig. 4L, S4I). RPL26L is a component of the ribosome 60S and may reflect a function of FAM111A in the nucleolus where FAM111A has been shown to localize outside of S phase (Tarnita et al., 2019). HMCES, on the other hand has recently been characterized as a suicide enzyme binding covalently to single strand DNA at abasic sites, forming DPC complexes which are degraded by the proteasome (Mohni et al., 2019). Supporting a role of FAM111A in HMCES degradation, FAM111A has recently been shown to degrade two proteins forming drug induced replication coupled DPCs, namely TOP1 and PARP adducts (Kojima et al., 2020). SPRTN, another PCNA binding protease, acts in parallel with the proteasome to degrade replication coupled DPCs (Larsen et al., 2019). Therefore, like SPRTN, FAM111A could be a parallel mechanism to degrade DPCs created by HMCES at abasic sites.

Collectively, our data show that FAM111A promotes origin activation and that unrestrained FAM111A peptidase activity is cytotoxic, in part due to its recruitment to PCNA (Fig. 4M). However, it remains unclear why patient mutations are promoting cell death. Our data suggest that in a disease context, FAM111A hyper degrades its substrates or targets novel substrates essential for fork stability. Moreover, these substrates, although in the vicinity of PCNA, may not be core replisome components but replisome accessory factors instead. Another possibility is that FAM111A is depleted in patient cells due to FAM111A enhanced autocleavage activity. Indeed, although partial FAM111A depletion was not cytotoxic in cancer cells, FAM111A may be essential during early stages of human development. Finally, our data suggest that FAM111A targets an inhibitor of origin firing, although no known regulators of origin firing qualified as FAM111A substrates in our assay. Instead, HMCES ranked as FAM111A top putative substrate. HMCES binds to abasic sites, one of the most common DNA lesions (Friedberg, 2006), and to ssDNA, which is extensively exposed upon origin firing. FAM111A could thus provide a mechanism to keep ssDNA HMCES-free upon S phase onset. However, as HMCES can be degraded by the proteasome, its accumulation alone in FAM111A depleted cells is probably insufficient to affect origin firing, suggesting that other FAM111A substrates remain to be identified. Nonetheless, HMCES is an interesting substrate as it suggests an unanticipated role for FAM111A in endogenous replication coupled DPCs. Overall, our data highlight how FAM111A may play positive roles in DNA replication in normal conditions while becoming harmful upon unrestrained expression of peptidase domain and patient mutations. Developing FAM111A peptidase domain inhibitors may thus be beneficial for our understanding of KCS2 and GCLEB syndrome’s etiology.

## Supporting information

Rios_Supplemental_Material

## ACKNOWLEDGEMENTS

We thank Julian Blow, Anja Groth, Karim Labib, John Rouse, Giulia Saredi, Jordan Taylor and all members of the Alabert lab for comments on the manuscript. We thank Anja Groth for the GFP-RPA1 and RFP-PCNA U-2-OS cell line, Karim Labib for the GINS1 antibody and John Rouse for the Flp-In T-Rex U-2-OS cell line. We thank the Centre for Advanced Scientific Technologies (CAST) for access to the FACS facility, the Proteomics and Mass Spectrometry Facility FingerPrints, the Dundee imaging facility, the Drug Discovery Unit for access to the Operetta high content microscope and the MRC PPU reagents and services facility. We would like to thank Manu Decker, Nina Svensen, Giulia Saredi and Federico Tiranelli for technical assistance. D.O.R.S. acknowledges support from MRC PhD studentship, V.A. and EGW received support from CRUK, S.B. received support from ERC. Research in Alabert lab is supported by CRUK Career Development fellowship C57404/A21782 and European Research Council ERC-STG715127. Research in Lamond lab is supported by Wellcome Trust Collaborative Award 206293/Z/17/Z. Research in Ponting lab is supported by the MRC MC_UU_00007/15.

## AUTHOR CONTRIBUTIONS

D.O.R.S and C.A. designed the project; D.O.R.S performed clonogenic assays, flow cytometry, immunoblotting, Cell Titer Glo assay, molecular DNA combing, IP-MS analysis and TMT-MS experiment. E.G.W. generated FAM111A inducible cell lines and performed the IP-MS and sub-cellular fractionation. D.O.R.S, V.A and S.B. performed NCC and HTM. LP performed the computational protein sequence analysis. H.J. performed the TMT Mass spectrometry analysis. D.O.R.S and C.A. wrote the manuscript with inputs from all authors.

