## Supplementary material for "FAM111A regulates replication origin activation and cell fitness": Rios_Supplemental_Material

[1]GRE, School of Life Sciences. University of Dundee DD15EH Dundee, UK

[2]MRC Human Genetics Unit, MRC Institute of Genetics and Molecular Medicine at the University of Edinburgh, Edinburgh EH4 2XU, UK

[3]GRE, School of Life Sciences. University of Dundee DD15EH Dundee, UK

### SUPPLEMENTAL MATERIAL

List of content:

Supplemental Material and Method

Figure S1, related to Figure 1

Figure S2, related to Figure 2

Figure S3, related to Figure 3

Figure S4, related to Figure 4

Figure S5, Uncropped immunoblots

Table S1, related to Figure 4

Table S2, related to Figure 4

### Supplemental materials and methods

**Cell lines and cell culture conditions.** U-2-OS (ATCC), HeLa S3 (ATCC), GFP-RPA1 and RFP-PCNA U-2-OS and Flp-In T-Rex U-2-OS cells were grown in DMEM (Gibco) containing 10% FBS, 1% penicillin/streptomycin and drugs for selection. Flag-HA-FAM111A-WT (WT)

and Flag-HA-FAM111B plasmids were generated in the pcDNA5/FRT/TO backbone. Flag-HA-Y24A-Y25A (PIPmt), Flag-HA-S541A, Flag-HA-R569H, Flag-HA-R569H-S541A, Flag-HA-Y511H, and Flag-HA-T338A plasmids were generated from Flag-HA-FAM111A-WT construct by site-directed mutagenesis. All plasmids were confirmed by Sanger sequencing. Cells inducible for FLAG-HA-FAM111A mutants and Flag-HA-FAM111B were generated in Flp-In T-REx U-2-OS cells by transfection of the above constructs with Lipofectamine 2000 (Invitrogen) according to the manufacturer's protocol, and selection with 100 μg/ml hygromycin.

**Transfections and siRNA.** siRNAs were introduced by Lipofectamine RNAiMAX (Invitrogen), according to manufacturer's recommendations. The following siRNAs were used. siFAM111A-1, GUAAUCAGUUUCAUGACACUAAAdAdG and siFAM111A-2, ACCUUGGUUUGAGAUACAUAAUGdAdA (Origene, SR324823); siFAM111A-p (pool), GCAUUGUGGGAGACGGAAU, UACUGAAACUGUCGGAAUA, CGAUUAAAGUAGUGAAACU, GGUCAAUGUGUAAGGGUGA [ON-TARGETplus SMART human FAM1111A (63091), Dharmacon, L-013926-01-0005]; siFAM111B-1: GCUUAAAGUGUCCAAUGAAAACTA (Origene, SR317776). Control siRNAs: siControl, CGUUAAUCGCGUAUAAUACGCGUdAdT, (Origene, SR30004), siControl-p (pool), UGGUUUACAUGUCGACUAA, UGGUUUACAUGUUGUGUGA, UGGUUUACAUGUUUUCUGA, UGGUUUACAUGUUUUCCUA (ON-TARGETplus Non-targeting Pool, Dharmacon, D-001810-10-15).

**Drug treatments.** Hydroxyurea: cells were treated with 3 mM hydroxyurea for 2 h for high throughput microscopy and immunoblotting or with 0.5 mM, 1 mM or 2 mM for 24 h for clonogenic assays. UCN-01: cells were treated with 300 nM for 2 h. Aphidicolin: 50 μg/ml aphidicolin for 2 h. Thymidine arrest: 2.2 mM for 17 h. Protein expression was induced in with 0.5-1 µg/ml tetracycline for 24 h.

**Sub-cellular fractionations.** Cytoplasmic, nuclear and chromatin-bound fractions were isolated as previously described (Mendez and Stillman, 2000). Briefly, to isolate chromatin, cells were resuspended ($4 \times 10^7$ cells/ml) in 0.1% Triton X-100 Buffer A (10 mM HEPES pH 7.9, 10 mM KCl, 1.5 mM $MgCl_2$, 0.34 M sucrose, 10% glycerol, 1 mM DTT, protease inhibitors) for 8 min on ice. Nuclei were pelleted at 1,300 g, 4 °C for 4 min (P1). The cytosolic supernatant (S1) was further clarified for 15 min at 20,000g, 4°C. P1 was washed in buffer A, and incubated with buffer B (3 mM EDTA, 0.2 mM EGTA, 1 mM DTT, protease inhibitors) for 30min on ice, centrifuged (4 min, 1,700g, 4°C), washed with buffer B and centrifuged again. For immunoprecipitations, the chromatin pellet (P3) was further resuspended in buffer C (20 mM HEPES pH7.9, 1.5 mM MgCl2, 420 mM NaCl, 25% glycerol, 1mM DTT, protease inhibitors) and incubated with benzonase (Millipore 70746-3) on ice for 30 min. For immunoprecipitation from whole cell extracts, cells were lysed with 420 mM NaCl, 20% glycerol, 10 mM HEPES pH 7.9, 0.1% NP-40, 1 mM EDTA, 1 mM DTT, protease inhibitors). Samples were then syringed with a 25G syringe 10 times and benzonase treated on ice for 30mins. To analyze soluble and chromatin-bound fractions by immunoblotting, cells were treated as previously described (Saredi et al., 2016). Briefly, cells were incubated in 0.5% Triton X-100 CSK buffer (10 mM PIPES pH 7, 100 mM NaCl, 300 mM sucrose, 3 mM MgCl2), supplemented with and protease and phosphatase inhibitors (5 mM sodium fluoride, 10 mM β-glycerolphosphate, 0.2 mM sodium vanadate, 10 µg/ml leupeptin, 10 µg/pepstatin, 0.1 mM PMSF) on ice for 10 min and centrifuged at 1,500g for 10 min to collect soluble proteins. Pellets were washed again in CSK buffer, resuspended with SDS lysis buffer (1% SDS, 50 mM Tris-HCl, pH 8.1, 10 mM EDTA, 1 mM PMSF, 10 mM $MgCl_2$, protease inhibitors) and treated with benzonase for 30 min. For immunoblotting of whole cell extracts, cells were lysed in SDS lysis buffer as above.

**Nascent Chromatin Capture.** NCC was performed as described previously (Alabert et al., 2014). Cells were synchronized by single thymidine block (2.2 mM) for 17 h and released into S phase with 24 µM 2'deoxycytidine for 3.5 h. Cells were labeled with 50 µM biotin-dUTP for 5 min in hypotonic buffer (50 mM KCl, 10 mM HEPES), supplemented with medium for 15 min, chased with biotin-dUTP free medium for indicated times and fixed for 15 min in 2% formaldehyde. Nuclei were isolated by douncing 20 times in sucrose buffer (0.3 M sucrose, 10 mM HEPES, pH 7.9, 1% Triton X-100, 2 mM MgOAc). Chromatin was solubilized in sonication buffer (10 mM HEPES pH 7.9, 100 mM NaCl, 2 mM EDTA, 1 mM EGTA, 0.2% SDS, 0.1% sodium sarkosyl, 1 mM PMSF) using Diagenoge Bioruptor (28 cycles, 30 sec on, 90 sec off, high intensity). Biotin-dUTP–labeled chromatin was purified on Streptavidin C1 Dynabeads (Invitrogen) overnight. Isolated nascent chromatin was boiled for 40 min in LSB (50 mM Tris-HCl at pH 6.8, 100 mM DTT, 2% SDS, 10% glycerol, Bromophenol blue).

**High-throughput microscopy.** U-2-OS cells were grown on clear bottom 96-well plates (Greiner) and either pre-extracted with cold 0.5% Triton X-100 CSK buffer for 5 min before fixation or directly fixed in 4% formaldehyde for 10 min. For EdU (5-ethynyl-2′-deoxyuridine) labelling, cells were incubated with 40 µM EdU for 20-30 min. EdU was detected using Click-iT™ Plus EdU Kit for Imaging (C10640). Plates were imaged on a Perkin Elmer Operetta high-content imaging system using a 20x objective. 35 fields per well were imaged, and approximately 2000 cells per condition were analyzed. Single cell fluorophore intensities were extracted using the Columbus system (Perkin Elmer). Cell cycle phases were gated based on

DAPI and EdU intensities. Graphs were generated using Tableau 2019.3 and GraphPad Prism.7 software.

**Clonogenic assay.** U-2-OS were transfected with siRNAs and after 24 h seeded in technical triplicates or duplicates of 2000 and/or 4000 cells in 10 cm dishes. 48 h after transfection, treatments were performed as indicated. Cells were then cultured in fresh medium for 10-14 days, fixed in methanol and stained with 25% Giemsa stain in methanol. Colony formation efficiency was determined blindly by manual colony counting and normalized to untreated controls. Each data point represents a technical replicate of 2000 cells seeded cells within each biological replicate.

**Cell proliferation assay.** U-2-OS were transfected with siRNAs and after 24 h seeded in clear bottom 96 well plates (Greiner) at 500 cells/well in triplicates. Cell viability was measured every 24 h for four days using CellTiter-Glo 2.0 Cell Viability Assay (Promega, G9242) according to manufacturer's recommendations.

**Molecular DNA combing.** Single-molecule analysis of DNA replication by molecular combing was performed as described in protocol 36 available from the EpiGeneSys Network of Excellence website. In brief, 48 h after siRNA transfections, U-2-OS cells were successively pulse labeled with 50 µM CldU and 250 µM IdU for 30 min each. Immediately after the pulse, cells were harvested, molded into low-melting agarose plugs, and DNA was extracted using Fiber Prep DNA extraction kit (EXT-002, Genomic Vision). DNA was combed on silanized coverslips (Genomic Vision) using FiberComb (Molecular Combing System, Genomic Vision) Combed coverslips were subsequently dehydrated at $60_{o}$C for 2h, denatured in denaturing solution (0.5 M NaOH, 1M NaCl) for 8 min, blocked with Block Aid solution (Invitrogen) for

30 min at 37 $_{o}$C, and probed with rat anti-BrdU (Abcam, ab6326, 1/25) and mouse anti-BrdU (BD Biosciences 347580, 2/25) antibodies for 1h, and anti-ssDNA (Merck Millipore MAB3034, 4/25) for 2 h. Coverslips were then washed with 0.05% PBST and incubated with anti-mouse Cy3.5 conjugated IgG (Abcam ab6946, 1/25) and anti-rat Cy5 conjugated IgG (Abcam ab6565, 1/25), and washed again. Immunolabeled coverslips were then dehydrated and sent for automated imaging using Genomic Vision EasyScan service. More than 300 fibers were analyzed for each condition. Measured distances were converted to kilobases by the constant stretching factor (1 µm = 2 kb). Inter-fork distances were determined based on CldU staining.

**Flow Cytometry.** For cell cycle analysis, cells were trypsinized, fixed with ice-cold 70% ethanol overnight at 4 $_{o}$C, treated with propidium iodide (PI) solution (50 µg/ml PI, 50 µg/ml RNaseA, and 0.1% Triton-X-100, 1% FBS in PBS) for 30 min and acquired using BD FACSCanto. Results were analyzed using FlowJo software.

**Immunoprecipitation from cell extracts.** Immunoprecipitations of endogenous FAM111A were performed with anti-FAM111A (Abcam, ab184572) or control IgG bound to Protein A Dynabeads (Thermo Fisher, 10002D), and incubated rotating overnight at 4 °C. Beads were s washed three times with wash buffer (20 mM HEPES pH 7.9, 150 mM KCl, 2.5 mM MgCl2, 1 mM DTT, 0.5 mM PMSF, 0.1% NP-40), boiled in LSB and subjected to SDS–PAGE separation on NuPAGE 4–12% gels (Invitrogen, NP0321).

**AP-MS analsysis**. Immunoprecipitated fractions were analyzed as follows. Samples were run on 4-12% NuPAGE gels (Invitrogen #NP0321) until top and bottom markers were separated by 1 cm. Gel slices were excised, broken into small pieces, successively washed for 15 min with 100 mM ammonium bicarbonate, 100 mM ammonium bicarbonate / acetonitrile (50:50) and acetonitrile, and dried *in vacuo*. Samples were then incubated in reducing solution (10 mM DTT, 20 mM ammonium bicarbonate) for 60 min at 56$_o$C, alkylated with 50 mM iodoacetamide in 20 mM ammonium bicarbonate for 30 min at room temperature, washed twice with 100 mM ammonium bicarbonate for 15 min and with acetonitrile for 10 min, and then dried *in vacuo*. Digestion was performed with trypsin (12.5 µg/ml) overnight at 30 °C. Samples were then extracted with consecutive acetonitrile washes, pooled and dried *in vacuo*. Label free peptide analysis was performed by the FingerPrints Proteomics Facility (University of Dundee) on a Q Exactive™ plus mass spectrometer (Thermo Scientific) coupled with a Dionex Ultimate 3000 Rapid Separation LC (Thermo Scientific). The following LC buffers were used: buffer A (0.1% formic acid in Milli-Q water (v/v)) and buffer B (80% acetonitrile and 0.1% formic acid in Milli-Q water (v/v). Aliquots of 10 µl were loaded at 10µl/min onto a trap column (100 µm × 2 cm, PepMap nanoViper C18 column, 5 µm, 100 Å, Thermo Scientific) equilibrated with 98% Buffer A. The trap column was washed for 5 min at the same flow rate and then switched in-line with PepMap RSLC C18 column (75 µm × 50 cm, 2 µm, 100 Å). The peptides were eluted from the column at 300 nl/min with a linear gradient of 2% -35% buffer B over 125 min, and then 98% buffer B by 127 min. The column was then washed with 98% buffer B for 20 min and re-equilibrated in 2% buffer B for 17 min. Q-Exactive plus was operated in positive mode using data dependent mode. A scan cycle comprised MS1 scan (m/z range from 330-1600, with a maximum ion injection time of 20 ms, a resolution of 70 000 and automatic gain control (AGC) value of 1x10$_6$ followed by 15 sequential dependent MS2 scans (with an isolation window set to 1.4 Da, resolution at 17500, maximum ion injection time at 100 ms and AGC 2x10$_5$. Dynamic exclusion was set at 45 s, stepped collision energy was set to 27 and fixed first mass to 100 m/z. Spectrum was acquired in centroid mode and unassigned charge states, charge states above 6 as well as singly charged species were rejected. The raw

AP-MS data were searched using MaxQuant (1.6.6.0) (Cox and Mann, 2008) and the Andromeda search engine software (Cox et al., 2011), and searched against *Homo Sapiens* database from Uniprot (SwissProt, December 2019). Data was searched with the following parameters: variable modification of Oxidation (M), deamidation (N, Q) and acetylation (protein N terminus) and fixed modification of Carbamidomethylation (C). MS/MS tolerance: FTMS was set at 10 ppm and ITMS at 0.06 Da. The FDR threshold was set to 1%, allowing for maximum peptide length of 8 and 2 missed cleavages. The ProteinGroups output table from MaxQuant was analyzed using Perseus (1.6.7.0) (Tyanova et al., 2016). Data was filtered to remove entries matched to "potential contaminant", "only identified by site" and "reverse". Entries with less than two unique peptides and not present in at least two FAM111A pull downs were also removed. Missing values were imputed from a normal distribution using default settings. Log2 ratios were compared using 2 sample Student's t-test.

**TMT MS analysis**. Cells were washed with ice-cold PBS three times, resuspended in urea lysis buffer (8 M urea in 100 mM Tris-HCl, pH 8, 1x Complete Mini EDTA-free Protease Inhibitor Cocktail), mixed for 15 min at room temperature and sonicated (Diagenode Bioruptor) for 30 cycles (30 s ON and 30 s OFF, high). The cell extracts were then cleared at 15000 g for 10 min, and supernatants were treated with 10 mM tris (2-carboxyethyl) phosphine (TCEP) at room temperature for 45 min, followed by 20 mM iodoacetamide at room temperature in the dark for 30 min. Protein samples were further processed using SP3 magnetic beads (GE Healthcare Life Sciences) to remove all the salts/contaminants and digested to peptides for TMT labelling (Hughes et al., 2019). In detail, samples were mixed with SP3 beads (1:10) in 70% acetonitrile and incubated for 10 min at room temperature. Beads were washed twice with 1 ml 70% ethanol, once with 1 mL acetonitrile, re-dissolved in 80 µL 50 mM ammonium bicarbonate, and digested with trypsin (1: 50, trypsin: protein) at 37 °C overnight.

Samples were acidified by adding 9 µL 10% formic acid and acetonitrile to 95%, incubating 10 min at room temperature. Beads were washed with acetonitrile and re-dissolved for in 2% DMSO. Equal amounts (100 µg) of each peptide sample were dried and dissolved in 100 µL 100 mM TEAB buffer. TMT labelling of each samples were followed by the TMT10plex™ Isobaric Mass Tag Labeling Kit (Thermo Fisher Scientific) manuals. TMT labeled peptides were fractionated using off-line high-pH RP chromatography. The samples were loaded onto XBridge BEH C18 Column (Waters, 130 Å, 3.5 µm, 4.6 mm x 250 mm), and separated on Dionex bioRS HPLC system. The gradient was as follows: solvents A (water), B (acetonitrile) and C (100 mM ammonium formate pH 9); 0-8 min, 5% B; 8-10 min, 5%-21.5% B; 10-21 min, 21.5%-48.8% B; 21-22min, 48.8%-90% B, 22-27 min, 90% B; 1 ml/min flow rate, with 10% C all through the gradient. Peptides were separated into 48 fractions, which were collected into 24 fractions. Fractions were subsequently dried and re-dissolved in 5% formic acid. NanoLC-MS/MS analysis of TMT labeled samples was performed on an Orbitrap Fusion Tribrid mass spectrometer (Thermo Fisher Scientific), coupled with a Dionex Ultimate 3000 RS nanoLC system (Thermo Fisher Scientific). Peptides were loaded on the trap column (75 µm × 2 cm PepMap-C18), using 0.1% TFA for 8 mins with 10 µL/min flow rate. The peptides were then eluted and separated by an EASY-Spray column (75 µm × 50 cm RP- C18), at a constant flow rate of 300 nl/min. The gradient was as follow: 0-8 min, 1%B; 8-15 min, 1%-10%B; 15-155 min, 10%-32%B; 155-165 min, 32%-75%B; 165-175 min, 75%-95%B; 175-180 min, 95%B; buffer A (0.1% formic in water (v/v)) and buffer B (80% acetonitrile and 0.1% formic acid in water (v/v)). The MS data were acquired by Xcalibur control software (v 4.1.31.9) and the TMT-SPS-MS3 method in top speed mode with cycle time 3 s. A scan cycle comprised MS1 scan from m/z range 375-1600, with a maximum ion injection time of 50 ms and automatic gain control (AGC) value of $4x10_5$, at a resolution of 120,000). The most intense ions were selected for fragmentation as MS2 using CID in the ion trap with 35% CID collision energy and an isolation window of 1.2 Th. The AGC target was set to 1.0 x $10^4$ with a maximum injection time of 50 ms and a dynamic exclusion of 60 s, the scan rate was set to ‘Turbo’. For accurate quantification of TMT peptides, a subsequent synchronous precursor selection (SPS)-MS3 scan was performed. Five MS2 fragment ions were selected using with a window of 2 Th and further fragmented using HCD collision energy of 65%. The MS3 fragments were then analyzed in the orbitrap with a resolution of 50,000, within m/z range 100-500. The AGC target was set to 5.0 x $10^4$ and the maximum injection time was set to 120 ms. All MS data of TMT fractions were analyzed together by MaxQuant (v 1.6.10.43) and searched against Homo Sapiens database from Uniprot (SwissProt, January 2020). The data was searched with the following parameters: fixed modification of carbamidomethyl (C), variable modifications of oxidation (M) and acetylation (protein N terminus), with maximum of 2 missed tryptic cleavages, reporter mass tolerance set to 0.03 ppm. The FDR threshold was set to 1% at Peptide Spectrum Match (PSM), peptides and protein levels. The TMT quantification was set to reporter ion MS3 type with 10plex TMT (LOT: UH285228). The ProteinGroups output table from MaxQuant was filtered to remove “Potential contaminant”, “Only identified by site” and “Reverse”. Proteins with less than 2 peptides and without unique peptides were also removed. After filtering, the ratio of FAM111AOE to control, and of siFAM111A to control were transformed to log2 ratio and normalized with a mean of 0, and factors with a standard deviation above 1 were removed. For GO-term analysis, the proteins further fulfilled the following criteria: the average intensity ratios for FAM111A OE to control and siFAM111A to control are > 0.6. For identification of putative FAM111A substrates, based on FAM111A ratio, the proteins further fulfil the following criteria: the mean ratio for FAM111A OE to control is < -1 and for siFAM111A to control is > 0.4.

**Western blotting and antibodies.** The following antibodies were used: FAM111A (Sigma HPA040176, 1:500-1:1000, Abcam ab184572, 1:500-1:1000), PCNA (Abcam ab29, clone PC10, 1:1000; ab18197, 1:1000 for immunofluorescence), H4K12ac (Millipore 07-595, 1/1000), Histone H3 (Abcam ab10799, 1:1000), MCM3 (Abcam ab4460, 1:1000), MCM2 (BD Bioscience 610701; 1:1000, Abcam ab4461 1:1000 for immunofluorescence), CDC45 (CST #11881, 1:1000), RPA2 (ab2175, clone 9H8, 1:1000, 1:300 for immunofluorescence), Histone H4 (Millipore 05-858, 1:1000), GINS1 (kind gift from Karim Labib's lab, 1:500), p-CHK1 S345 (CST #2348, 1:1000) CHK1 (Santa Cruz sc-56291, 1:1000, p-CHK2 T68 (CST #2661, 1:1000) p-RPA S4/S8 (Bethyl A300-245A-M 1:1000), p-H2A.X S139 (γH2A.X) (CST #2577, 1:1000, 1:500 for immunofluorescence), p-RPA S33 (Bethyl A300-246A, 1:1000), anti-HA (CST #3724, 1:1000; Biolegend 901501, 1:000 for immunofluorescence) , GAPDH (CST #2118, 1:1000), AIF (CST #5318, 1:1000), Vimentin (CST #57411:1000), MEK1/2 (CST #8727, 1:1000), FAM111B (Invitrogen PA5-58474, 1:1000-1:2000). Secondary antibodies conjugated with horseradish peroxidase (HRP) were from Jackson ImmunoResearch Labs. Fluorescent dye conjugated antibodies were sourced from Li-Cor. Signals were revealed by chemiluminescence substrate from Pierce (SuperSignal West Pico or SuperSignal West Femto) and imaged using ChemiDoc XRS+ or Licor Odyssey imaging systems.

**Gene ontology (GO) analysis.** GO enrichment analysis including biological process (BP), Kyoto Encyclopedia of Genes and Genomes (KEGG) pathway and Biocarta enrichment analysis for TMT based mass spectrometry data set was performed using DAVID functional annotation tool (ver. 6.8). All proteins identified were used as background.

**Computational protein sequence analysis for identification of FAM111A UBL repeats.**

Multiple sequence alignments were generated with T-Coffee using default parameters (Notredame et al., 2000), slightly refined manually and visualized with the Belvu program (Sonnhammer and Hollich, 2005). Sequences were named according to their UniProt identifiers (Wu et al., 2006). Profiles of the alignment as global hidden Markov models (HMMs) were generated using HMMer (Eddy, 1996; Finn et al., 2011). Profile-based sequence searches were performed against the Uniref50 protein sequence database (Wu et al., 2006) using HMMsearch (Eddy, 1996; Finn et al., 2011). Remote homology analyses were performed using HHpred profile-to-profile comparisons (Soding et al., 2005). Profile-to-profile (HHpred) matches were evaluated in terms of an E-value, which is the expected number of non-homologous proteins with a score higher than that obtained for the database match. An E-value much lower than one indicates statistical significance. Secondary structure predictions were performed using PsiPred (Jones, 1999). The final UBL alignment was obtained using a combination of profile-to-profile comparisons (Soding et al., 2005) and sequence alignments derived from structural super-positions of a selection of UBL domains whose tertiary structure is known (PDB IDs: 1WFY, 5J2R, 2D07 and 3EEC) (Holm and Sander, 1995). Figures were generated using Inkscape (http://inkscape.org/). Structures and 3D models were analyzed using Pymol (http://www.pymol.org). Structural models were created using Modeller (Sali and Blundell, 1993). Surface electrostatic potential representations were generated using Pymol APBS (Adaptive Poisson-Boltzmann Solver) interface and were colored according to charge levels ranging from -5 kT/e (red) to +5 kT/e (blue).

**Statistical Analysis.** For statistical analysis, unpaired Student's t-tests and Mann Whitney tests were performed using Prism.7. P-values are indicated by asterisks (P < 0.001 [***], P < 0.01 [**], and P < 0.05 [*]), and ns indicates nonsignificant.

**Supplementary figure legends**

**Figure S1. A.** Inputs from Fig. 1B. **B.** Cell cycle progression from Fig. 1C analyzed by flow cytometry based on propidium iodide. **C.** Inputs from Fig. 1C. **D.** Immunoblot of whole cell extracts from siRNA-transfected U-2-OS cells for 48 h. **E.** HTM gating strategy for cell cycle analysis, based on EdU and DAPI cell intensities. **F**. Cell cycle distribution of U-2-OS siRNA-transfected cells. Data are represented as mean and SD of three independent experiments. **G.** Clonogenic survival assay of U-2-OS siRNA-transfected cells. Data are represented as colony numbers relative to control siRNA from all technical replicates in four independent experiments. **H.** Cell proliferation of U-2-OS siRNA-transfected cells measured by Cell Titer Glo 2.0 assay. Data a represented as mean and SD of three technical replicates; n=2. **I.** Inter-fork distances 48h after siRNA treatments. Top, inter-fork distance measurement schematic. Bottom, distribution of inter CldU track length. Red bar represents the median; n > 100. **J.** Size distribution of CldU track length. Cells were treated with 300 nM UCN-01 for 1 h prior to and during labelling as in Fig. 1E. Red bar represents the median; n > 400. **K.** Distribution of inter fork distances (IFD) as in I. Red bar represents the median. n > 100. **L**. Induction of origin firing upon UCN-01 treatment measured by DNA combing. Data are shown as 1/($IFD_{UCN-01}$/$IFD_{untreated}$). **M.** Cell cycle progression from Fig. 1I of cells synchronized with thymidine, released into S phase for 5 h and analyzed by flow cytometry based on propidium iodide staining. **N.** Immunoblot of chromatin fractions from asynchronous siRNA transfected U-2-OS cells pretreated with 300 nM UCN-01 for 2 h. **O**. HTM gating strategy used to determine S phase cells based on chromatin bound PCNA and DAPI intensities. **P.** Nuclear RPA2 intensities detected by HTM in directly fixed cells. n > 2500. Data are representative of two (A-C, N) three (D-M, P), and one (I-L) independent experiments. G-H, P unpaired Student's t-test. I-K, Mann-Whitney test. ***P < 0.001, ** P < 0.01, *P<0.05.

**Figure S2. A.** Schematic representation of active fork and dormant origin in unchallenged conditions and upon HU treatment. In response to HU, ongoing forks stall and dormant origins fire, leading to increased amount of ssDNA exposed and subsequent RPA loading. **B**. Chromatin binding of GFP-RPA1 in pool siRNA transfected cells. Blue, PCNA positive cells, grey, PCNA negative cells. **C.** Quantification of chromatin bound RPA2 in S phase in pool siRNA transfected U-2-OS cells. **D.** Quantification of chromatin bound GFP-RPA1 in S phase in siRNA transfected cells. **E**. Immunoblot of whole cell extracts from pool siRNA-transfected cells. For B-E, cells were treated with 3 mM HU for 2 h. For C and D, S phase cells were gated based on chromatin bound PCNA intensities. Data are representative of three (B, C, D) and two (E) independent experiments.

**Figure S3. A.** Immunoblot of whole cell extracts 24 h after induction of FAM111A WT, PIPmt or S541A with either 0.5 µg/ml (+) or 1.0 µg/ml (++) tetracycline (tet). **B**. Chromatin binding of γH2A.X shown as a function of DAPI intensities. Line indicates threshold to determine γH2A.X positive cells. Colors represent cell cycle stage gated as shown in Fig. S3C. **C.** Gating strategy based on EdU and DAPI intensities used in Fig. 3D, G-H, and S3B, D to determine cell cycle stage. eS, mS and lS correspond to early, mid and late S phase, respectively. **D**. Chromatin binding of Flag-HA-FAM111A shown as a function of DAPI intensities. Line indicates threshold to determine FAM111A positive cells. **E**. Gating strategy for quantification of mid S phase EdU negative cells in Fig. 3E. For illustrative purposes, data represent 0.5 µg/ml tet induction of Flag-HA-FAM111A overexpression. **F**. Conservation of FAM111A and FAM111B paralogues in mammals. FAM111 paralogues groups are indicated by colored bars to the left of the alignment: FAM111A and FAM111B subfamilies are indicated in red and yellow, respectively. Sequences are named according to their species name followed by their

UniProt identifier. Conserved domains in the FAM111 protein family are highlighted in different colors: UBL repeats 1 (labelled U1) and 2 (U2), and the Trypsin-like domain are colored in red, yellow, and green, respectively. The PIP box is labeled in blue. The ssDNA binding region is shown in red, with borders marked by *. The positions of known disease related missense mutations in human FAM111A are labeled (black rectangles). The positions of the catalytic triad in the Trypsin-like domain are also labeled (green rectangles). The position putatively related with ssDNA binding (F231, orange rectangle) and the autocleavage site (N-terminal to G335, violet rectangle) are labelled. The amino acid coloring scheme indicates the average BLOSUM62 score (correlated to amino acid conservation) in each alignment column: black (greater than 3), grey (between 3 and 1.5) and light grey (between 1.5 and 0.5). **G**. Immunoblot of whole cell extracts 24 h after induction of FAM111A WT, Y511H, T338A, or R569H with either 0.5 µg/ml (+) or 1.0 µg/ml (++) tetracycline (tet). Data are representative three (A, B, D) or two (G) independent experiments.

**Figure S4. A.** Schematic of cell fractionation strategies. **B**. Immunoblot of distinct cell fractions prepared as described in A. **C**. Immunoblot of endogenous FAM111A immunoprecipitation from chromatin fractions and whole cell extracts. IP, immunoprecipitation; UB, unbound fraction. **D**. Inputs from NCC analysis in Fig. 4D. **E**. Analysis of chromatin bound RPA in siRNA transfected U-2-OS cells after treating with 3 mM HU for 2 h. n= 685, 490, 532, 466, 534. **F.** Immunoblot of whole cell extracts from siRNA-transfected U-2-OS cells. **G**. GO-term analysis for biological pathways. Factors from upregulated in either FAM111A overexpression (OE) or FAM111A knock down (siFAM111A) compared to control cells, with log2(ratio) > 0.6, were selected for DAVID based analysis. Control represents all factors identified by TMT mass spectrometry. **H**. Replisome components identified by TMT mass spectrometry, shown as in Fig. 4L. PCNA was the most depleted upon FAM111A overexpression and enriched upon FAM111A depletion. **I**. Top putative FAM111A substrates. Data are representative of three (B, C) and two (D-F, H, I) independent experiments.

**Table S1. FAM111A interactome.** AP-MS analysis of FAM111A in whole cell extract and chromatin fraction in U-2-OS cells.

**Table S2. FAM111A putative substrates.** TMT based mass spectrometry analysis of whole cell extracts of Flag-HA FAM111A inducible U-2-OS cells 48 h after siRNA transfection and 24 h after 1.0 µg/mL tetracycline induction.

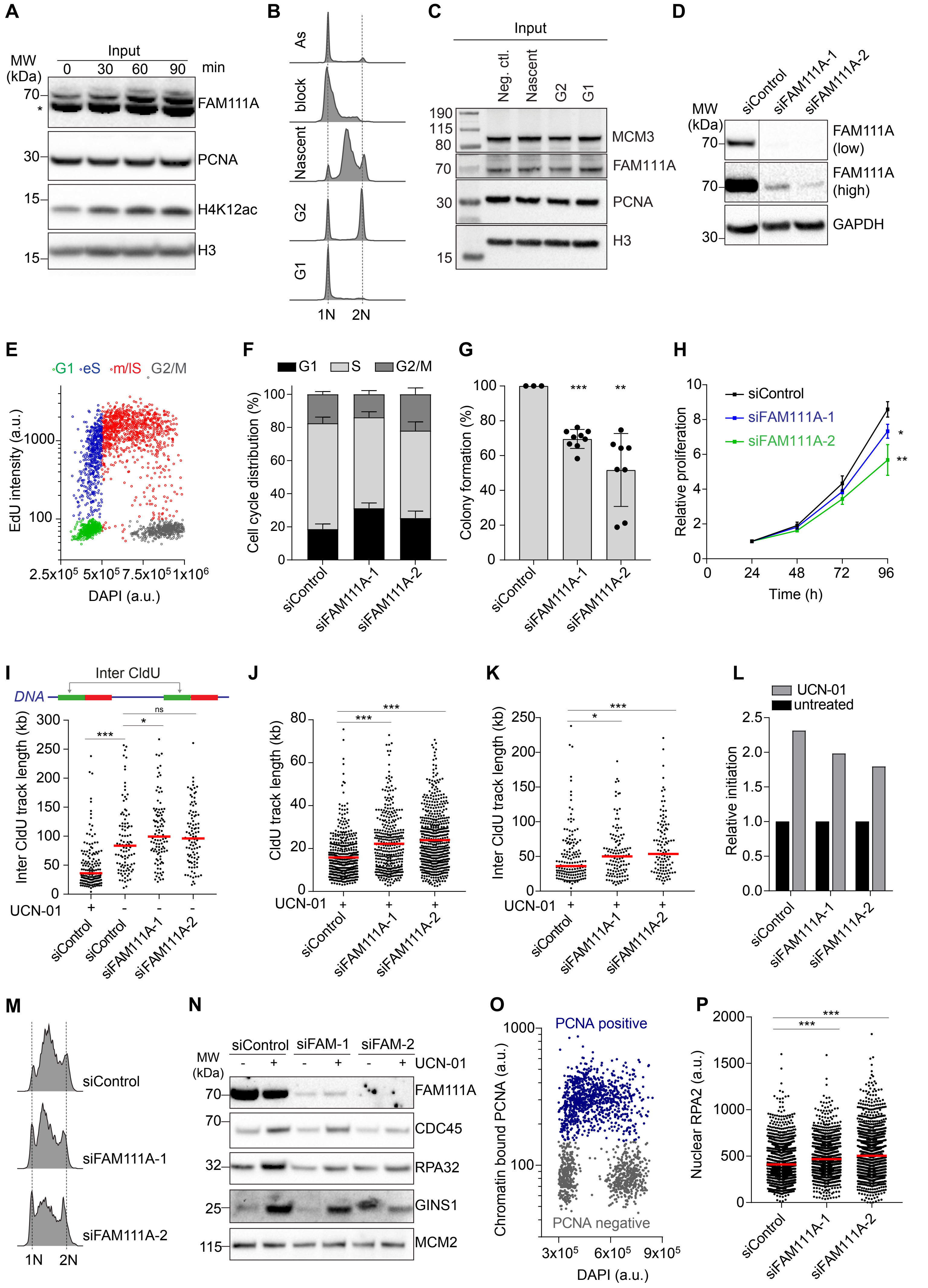
A
MW (kDa)
Input
0 30 60 90 min
FAM111A
PCNA
H4K12ac
H3
B
As
block
Nascent
G2
G1
1N 2N
C
Input
Neg. ctl.
Nascent
G2
G1
MCM3
FAM111A
PCNA
H3
D
siControl
siFAM111A-1
siFAM111A-2
FAM111A (low)
FAM111A (high)
GAPDH
E
G1 eS m/lS G2/M
EdU intensity (a.u.)
DAPI (a.u.)
F
G1 S G2/M
Cell cycle distribution (%)
G
Colony formation (%)
H
siControl
siFAM111A-1
siFAM111A-2
Relative proliferation
Time (h)
I
Inter CldU
DNA
Inter CldU track length (kb)
UCN-01
J
CldU track length (kb)
K
Inter CldU track length (kb)
L
UCN-01
untreated
Relative initiation
M
siControl
siFAM111A-1
siFAM111A-2
1N 2N
N
siControl siFAM-1 siFAM-2
UCN-01
FAM111A
CDC45
RPA32
GINS1
MCM2
O
PCNA positive
PCNA negative
Chromatin bound PCNA (a.u.)
DAPI (a.u.)
P
Nuclear RPA2 (a.u.)

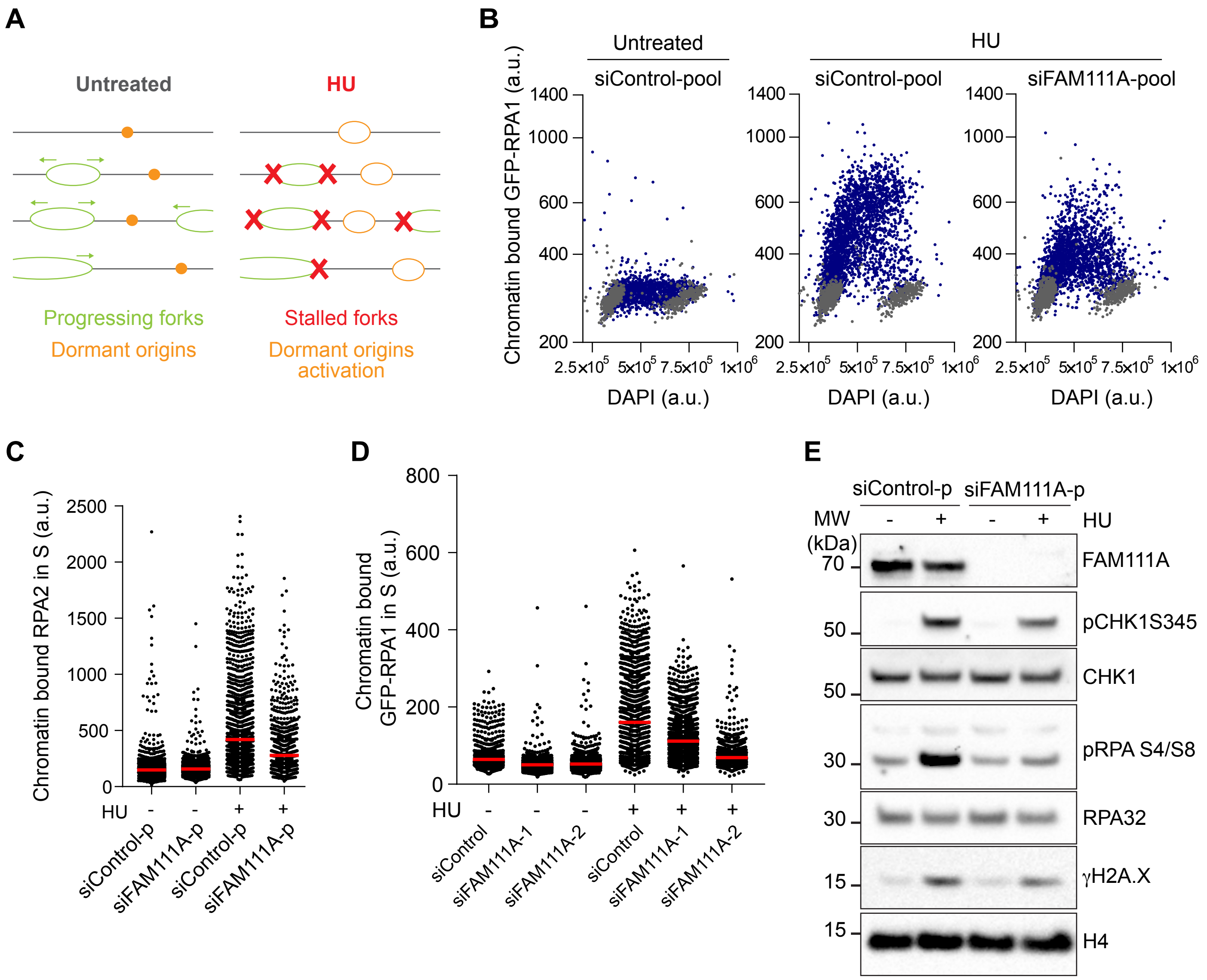
A
Untreated
HU
Progressing forks
Dormant origins
Stalled forks
Dormant origins activation
B
Untreated
HU
siControl-pool
siControl-pool
siFAM111A-pool
Chromatin bound GFP-RPA1 (a.u.)
DAPI (a.u.)
C
Chromatin bound RPA2 in S (a.u.)
HU
siControl-p
siFAM111A-p
siControl-p
siFAM111A-p
D
Chromatin bound GFP-RPA1 in S (a.u.)
HU
siControl
siFAM111A-1
siFAM111A-2
siControl
siFAM111A-1
siFAM111A-2
E
siControl-p
siFAM111A-p
MW (kDa)
HU
FAM111A
pCHK1S345
CHK1
pRPA S4/S8
RPA32
γH2A.X
H4

RIOS_FigS3

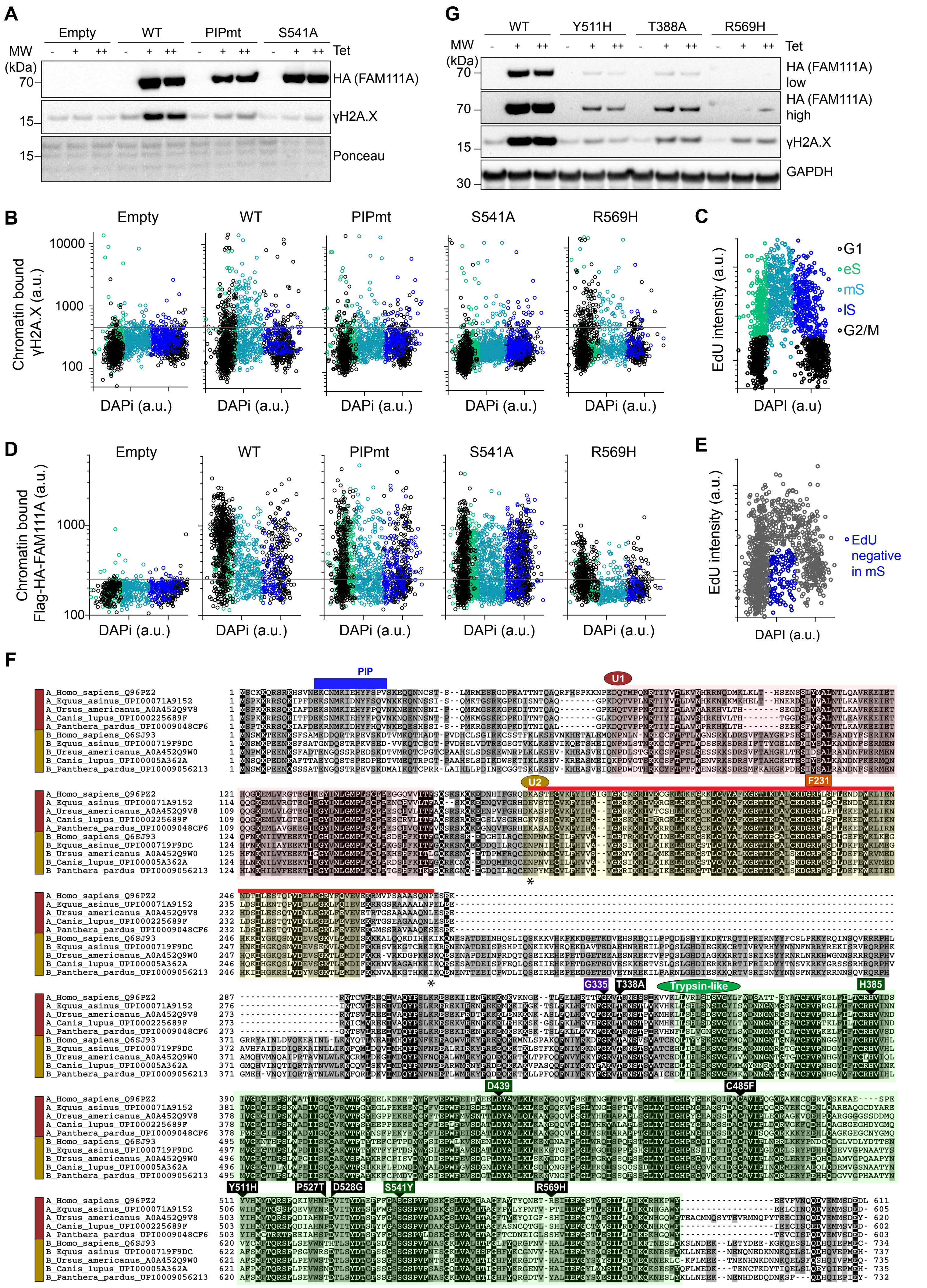
A
Empty
WT
PIPmt
S541A
MW (kDa)
Tet
- + ++
70
15
HA (FAM111A)
γH2A.X
Ponceau
G
WT
Y511H
T388A
R569H
HA (FAM111A) low
HA (FAM111A) high
γH2A.X
GAPDH
30
B
Empty
WT
PIPmt
S541A
R569H
Chromatin bound γH2A.X (a.u.)
10000
1000
100
DAPi (a.u.)
C
EdU intensity (a.u.)
DAPI (a.u)
G1
eS
mS
lS
G2/M
D
Chromatin bound Flag-HA-FAM111A (a.u.)
E
EdU negative in mS
DAPI (a.u.)
F
PIP
U1
U2
F231
G335
T338A
Trypsin-like
H385
D439
C485F
Y511H
P527T
D528G
S541Y
R569H
A_Homo_sapiens_Q96PZ2
A_Equus_asinus_UPI00071A9152
A_Ursus_americanus_A0A452Q9V8
A_Canis_lupus_UPI000225689F
A_Panthera_pardus_UPI0009048CF6
B_Homo_sapiens_Q6SJ93
B_Equus_asinus_UPI000719F9DC
B_Ursus_americanus_A0A452Q9W0
B_Canis_lupus_UPI00005A362A
B_Panthera_pardus_UPI0009056213
611
605
620
602
603
734
737
735
735
732

RIOS_FigS4

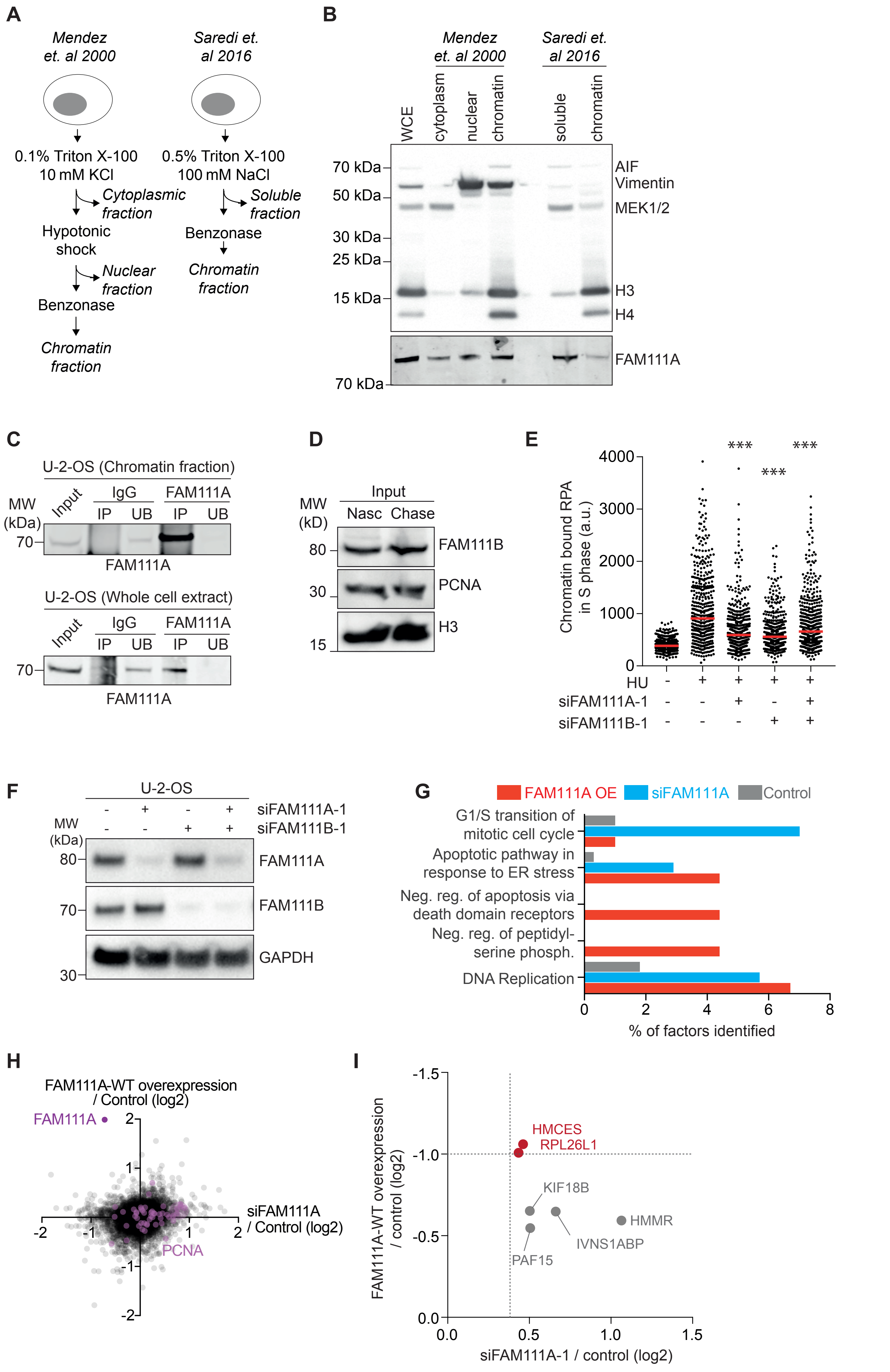
A
Mendez et. al 2000
Saredi et. al 2016
0.1% Triton X-100
10 mM KCl
Cytoplasmic fraction
Hypotonic shock
Nuclear fraction
Benzonase
Chromatin fraction
0.5% Triton X-100
100 mM NaCl
Soluble fraction
Benzonase
Chromatin fraction
B
WCE
cytoplasm
nuclear
chromatin
soluble
chromatin
70 kDa
50 kDa
30 kDa
25 kDa
15 kDa
70 kDa
AIF
Vimentin
MEK1/2
H3
H4
FAM111A
C
U-2-OS (Chromatin fraction)
MW (kDa)
Input
IgG
FAM111A
IP
UB
IP
UB
70
FAM111A
U-2-OS (Whole cell extract)
70
FAM111A
D
MW (kD)
Input
Nasc
Chase
80
30
15
FAM111B
PCNA
H3
E
Chromatin bound RPA in S phase (a.u.)
4000
3000
2000
1000
0
***
***
***
HU
siFAM111A-1
siFAM111B-1
F
U-2-OS
siFAM111A-1
siFAM111B-1
MW (kDa)
80
70
30
FAM111A
FAM111B
GAPDH
G
FAM111A OE
siFAM111A
Control
G1/S transition of mitotic cell cycle
Apoptotic pathway in response to ER stress
Neg. reg. of apoptosis via death domain receptors
Neg. reg. of peptidyl-serine phosph.
DNA Replication
0
2
4
6
8
% of factors identified
H
FAM111A-WT overexpression / Control (log2)
FAM111A
PCNA
siFAM111A / Control (log2)
I
FAM111A-WT overexpression / control (log2)
-1.5
-1.0
-0.5
0.0
HMCES
RPL26L1
KIF18B
HMMR
IVNS1ABP
PAF15
0.0
0.5
1.0
1.5
siFAM111A-1 / control (log2)

RIOS_FigS5

**Fig. 1A**

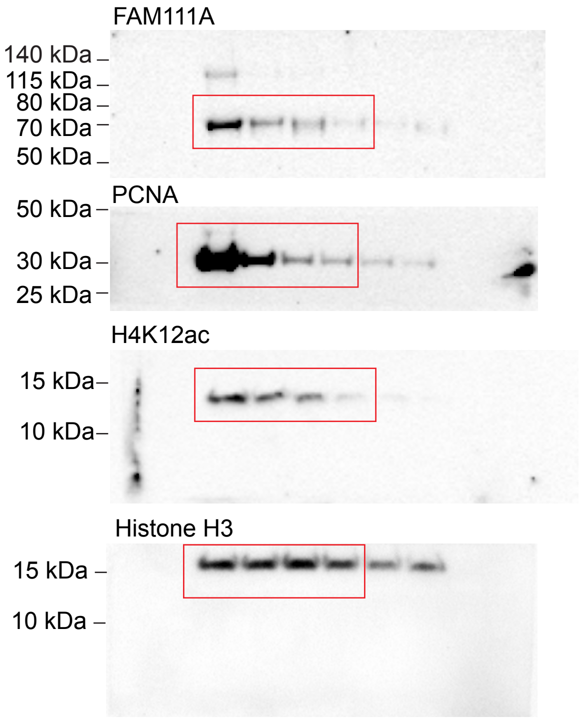


**Fig. 1B**

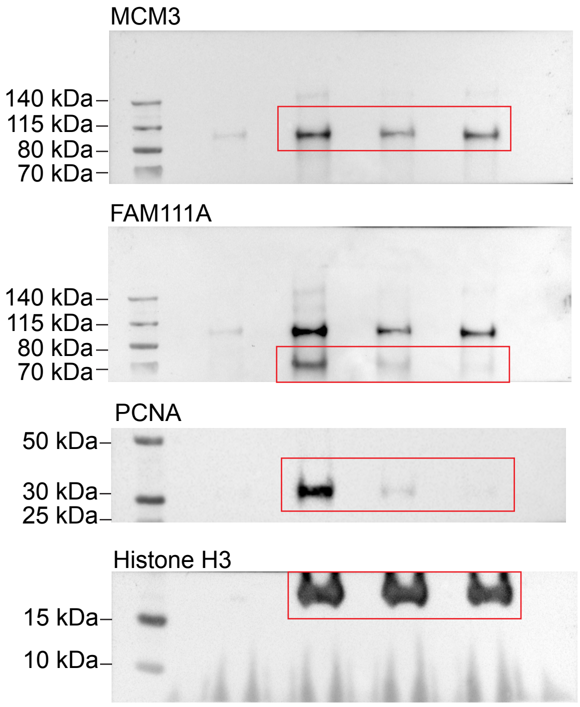


**Fig. 1I**

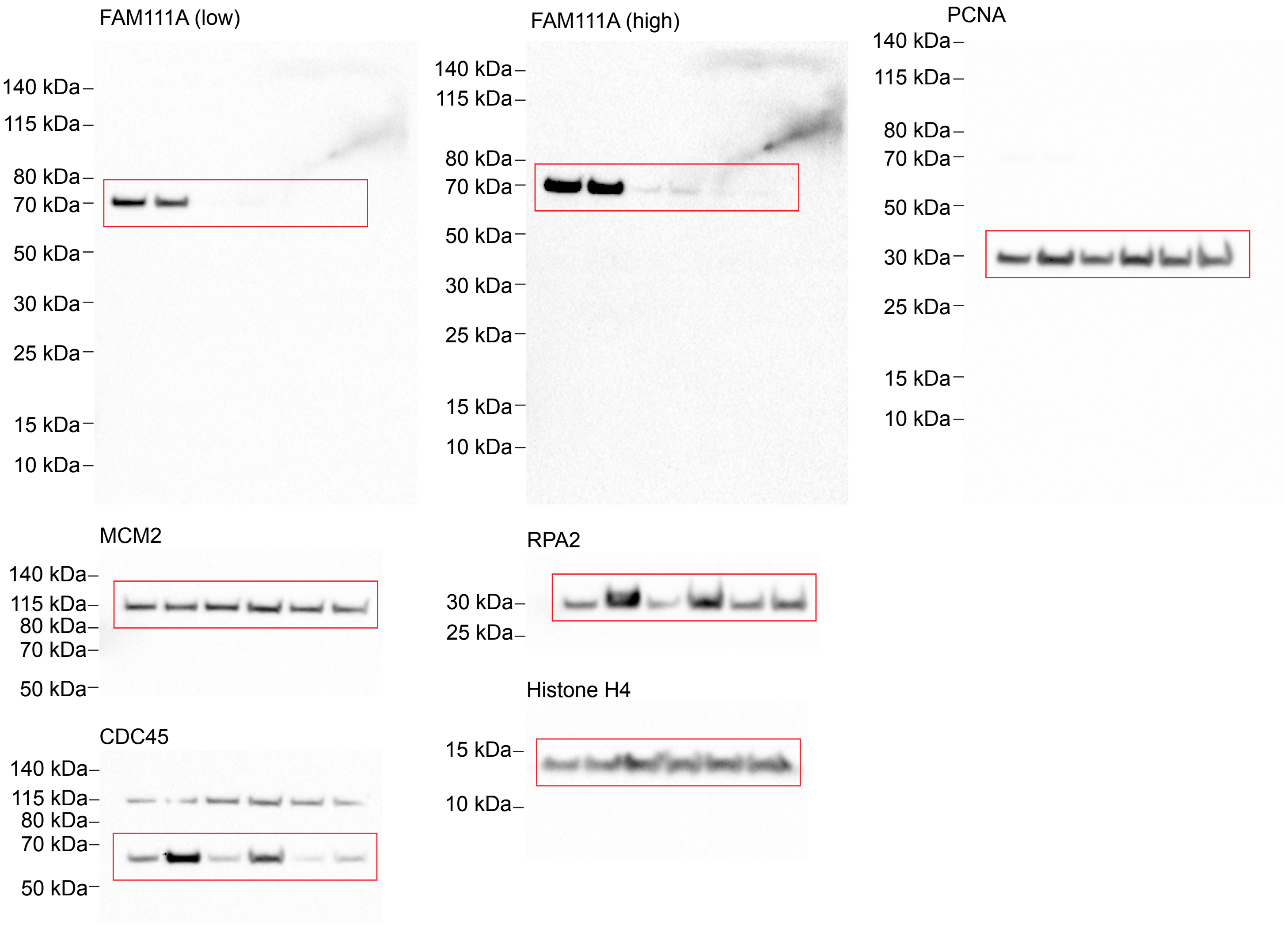

### Fig. 1K

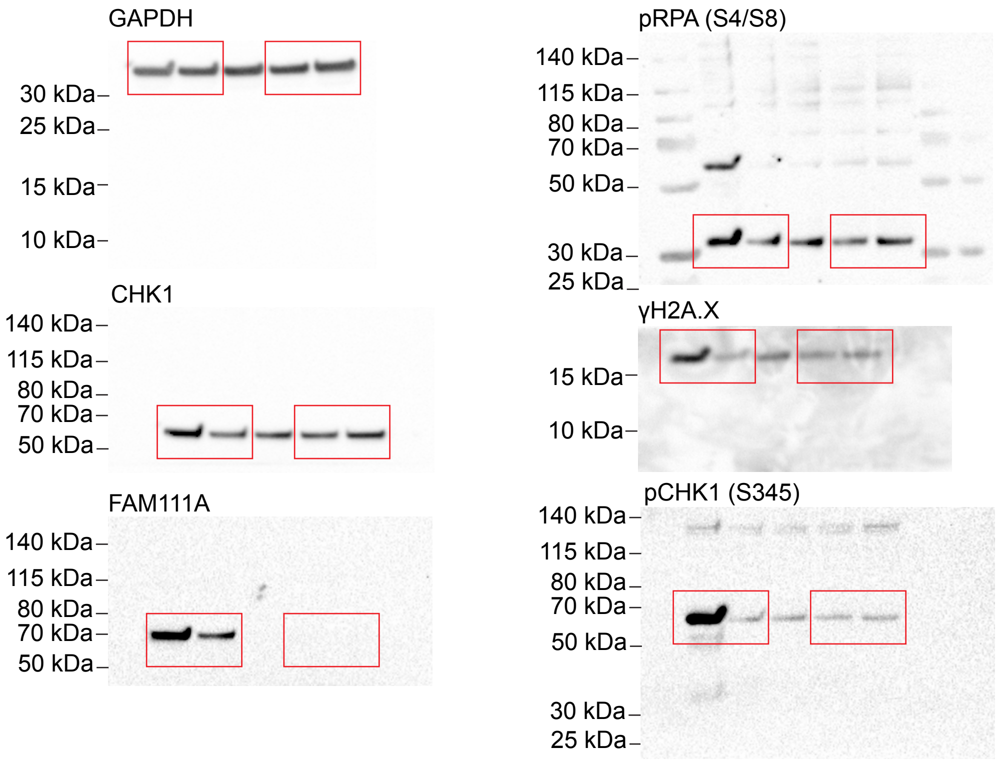
GAPDH
30 kDa
25 kDa
15 kDa
10 kDa
CHK1
140 kDa
115 kDa
80 kDa
70 kDa
50 kDa
FAM111A
140 kDa
115 kDa
80 kDa
70 kDa
50 kDa
pRPA (S4/S8)
140 kDa
115 kDa
80 kDa
70 kDa
50 kDa
30 kDa
25 kDa
γH2A.X
15 kDa
10 kDa
pCHK1 (S345)
140 kDa
115 kDa
80 kDa
70 kDa
50 kDa
30 kDa
25 kDa

### Fig. 2F

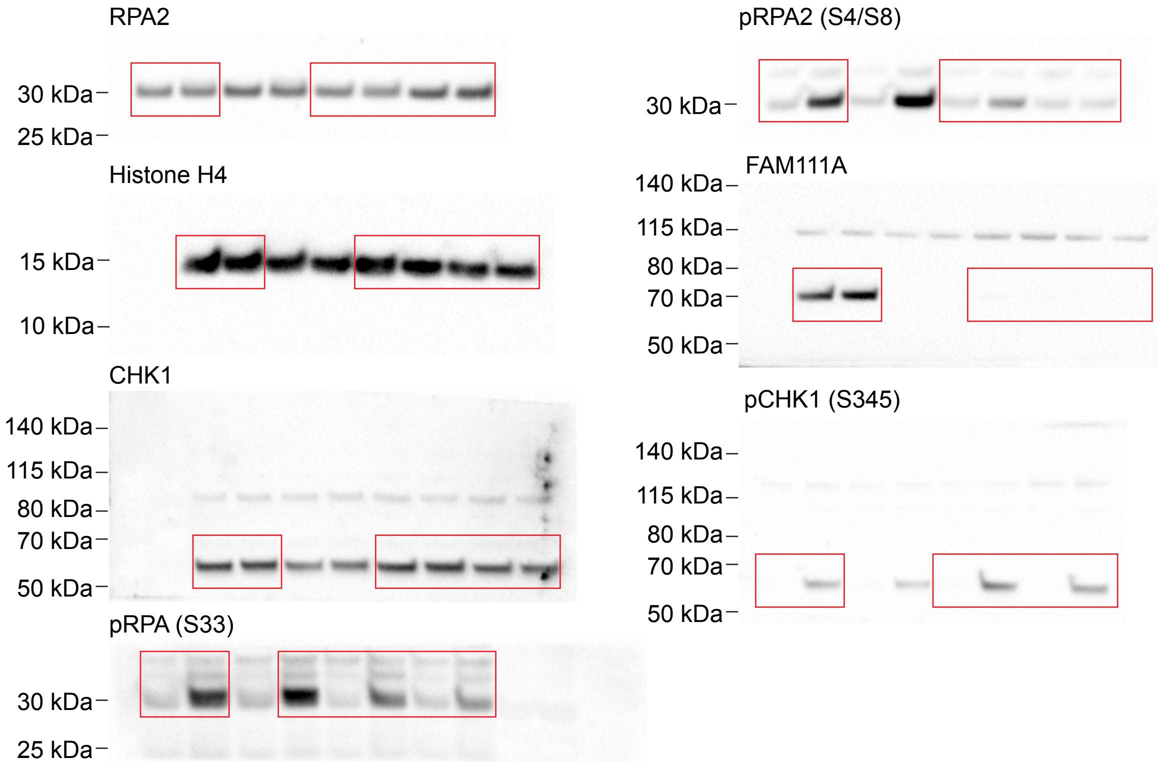
RPA2
30 kDa
25 kDa
Histone H4
15 kDa
10 kDa
CHK1
140 kDa
115 kDa
80 kDa
70 kDa
50 kDa
pRPA (S33)
30 kDa
25 kDa
pRPA2 (S4/S8)
30 kDa
FAM111A
140 kDa
115 kDa
80 kDa
70 kDa
50 kDa
pCHK1 (S345)
140 kDa
115 kDa
80 kDa
70 kDa
50 kDa

### Fig. 3J

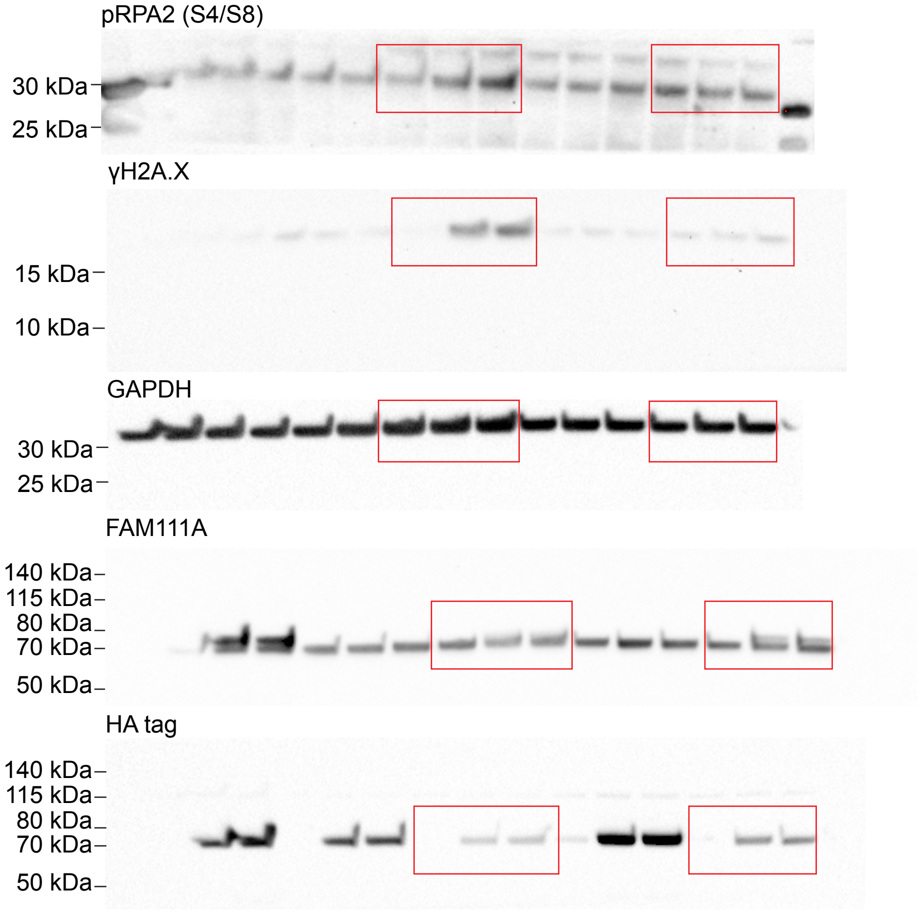


### Fig. 4D

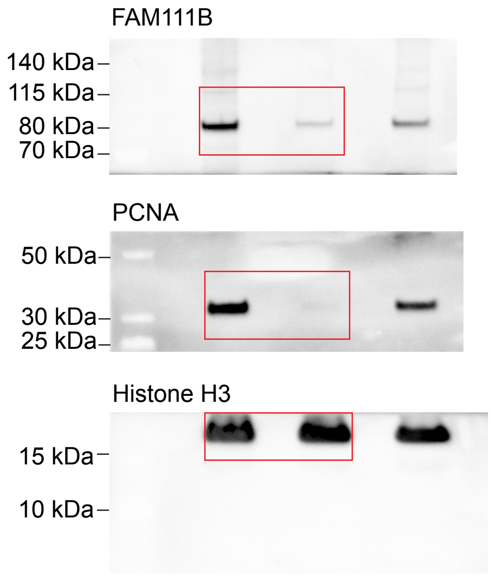


### Fig. 4F

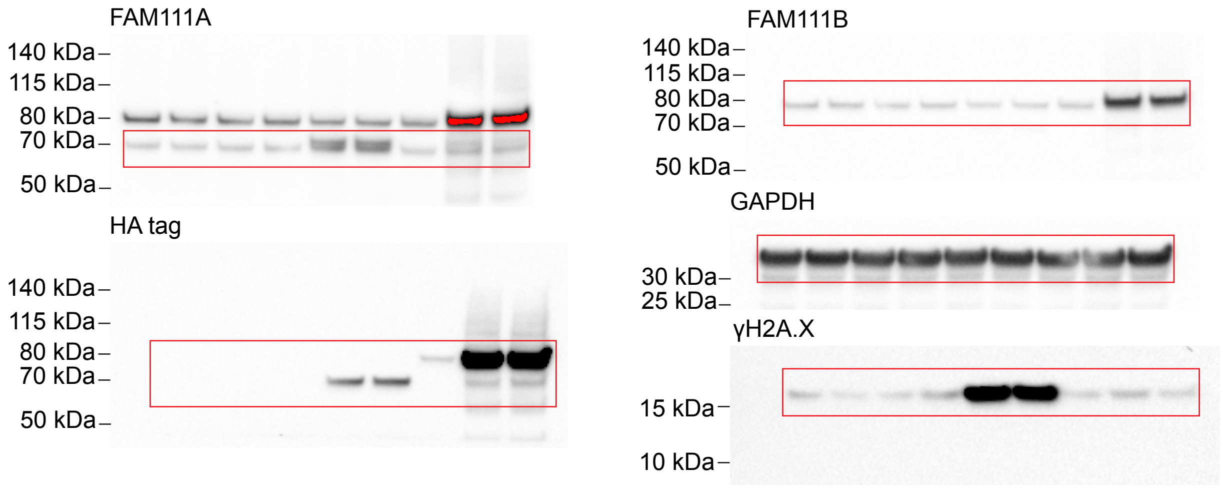


### Fig. 4K

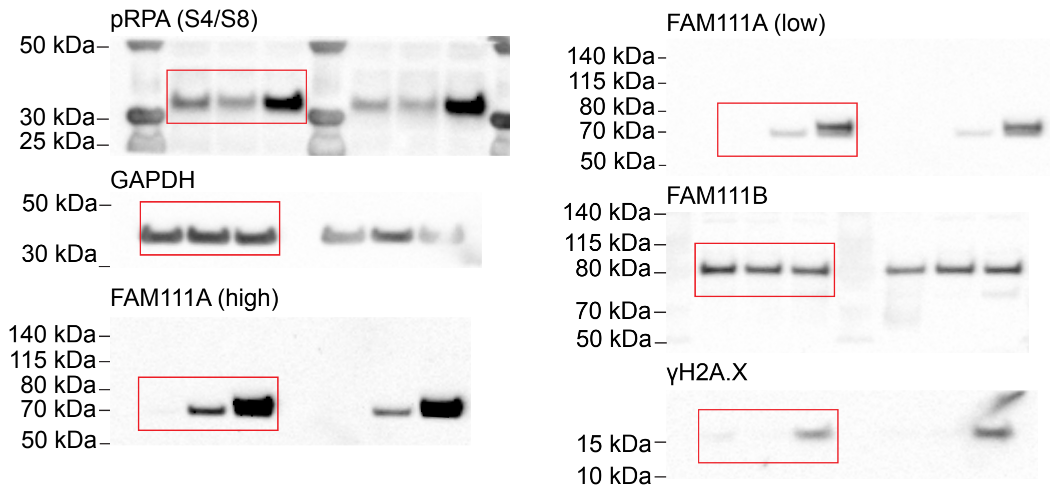

RIOS_FigS5

**Fig. S1A**

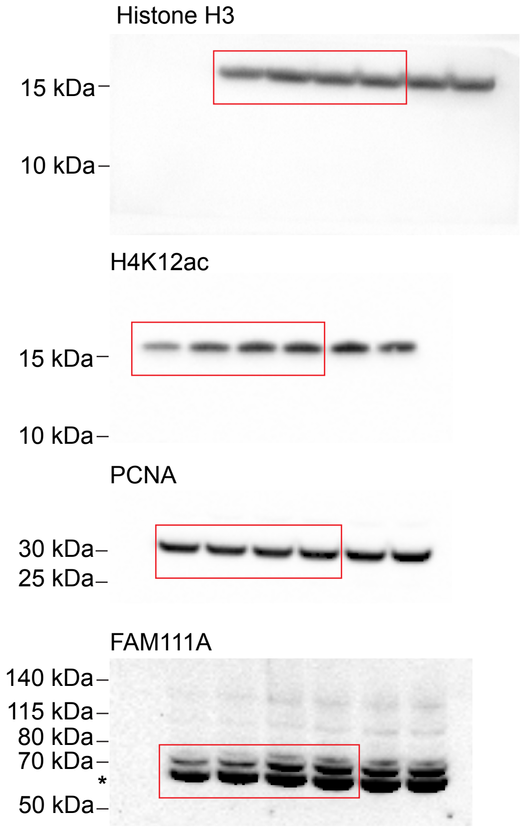


**Fig. S1C**

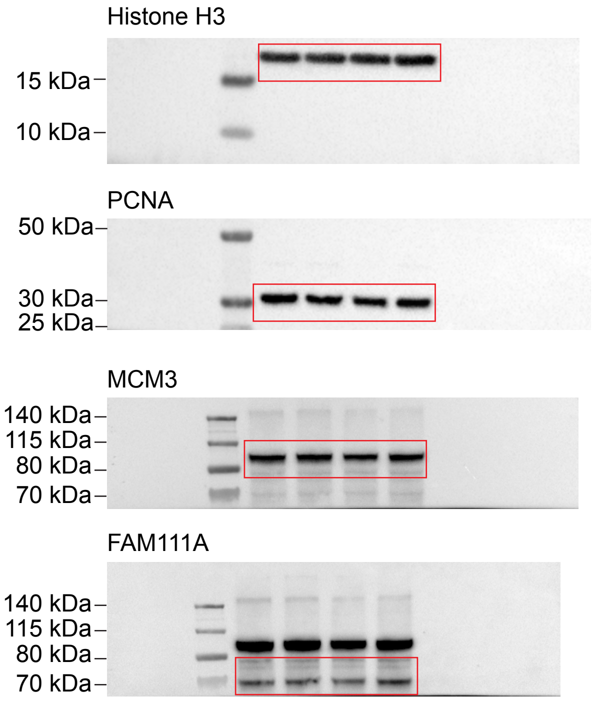


**Fig. S1D**

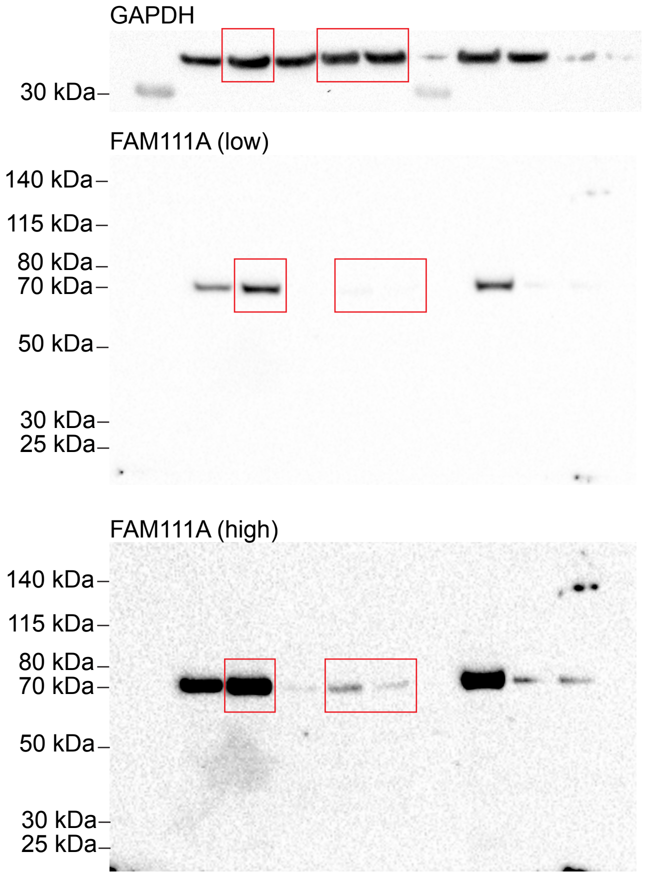


**Fig. S1N**

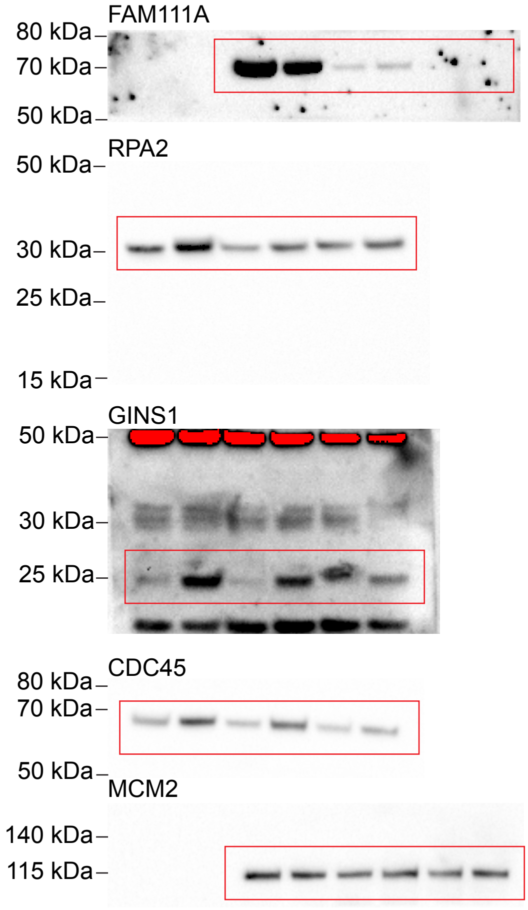

**Fig. S2E**

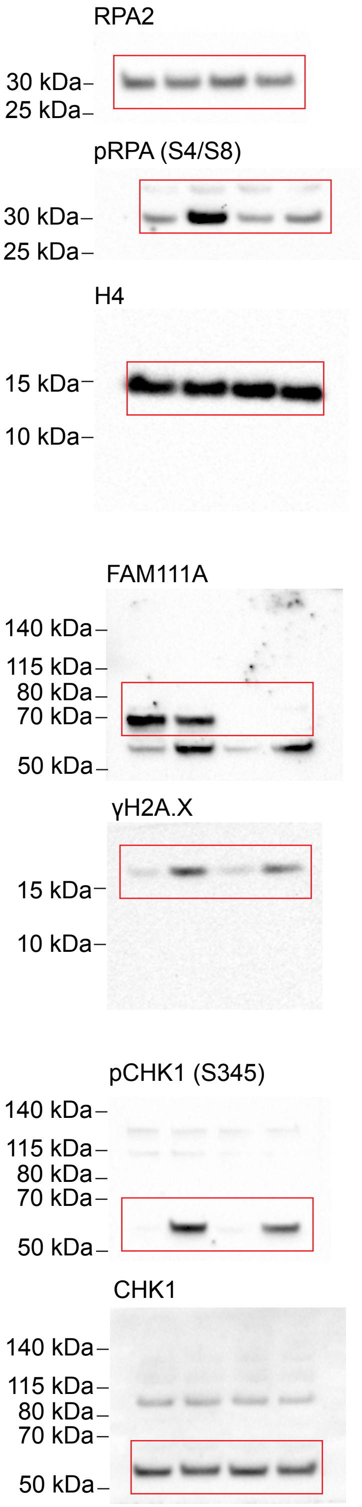


**Fig. S3A**

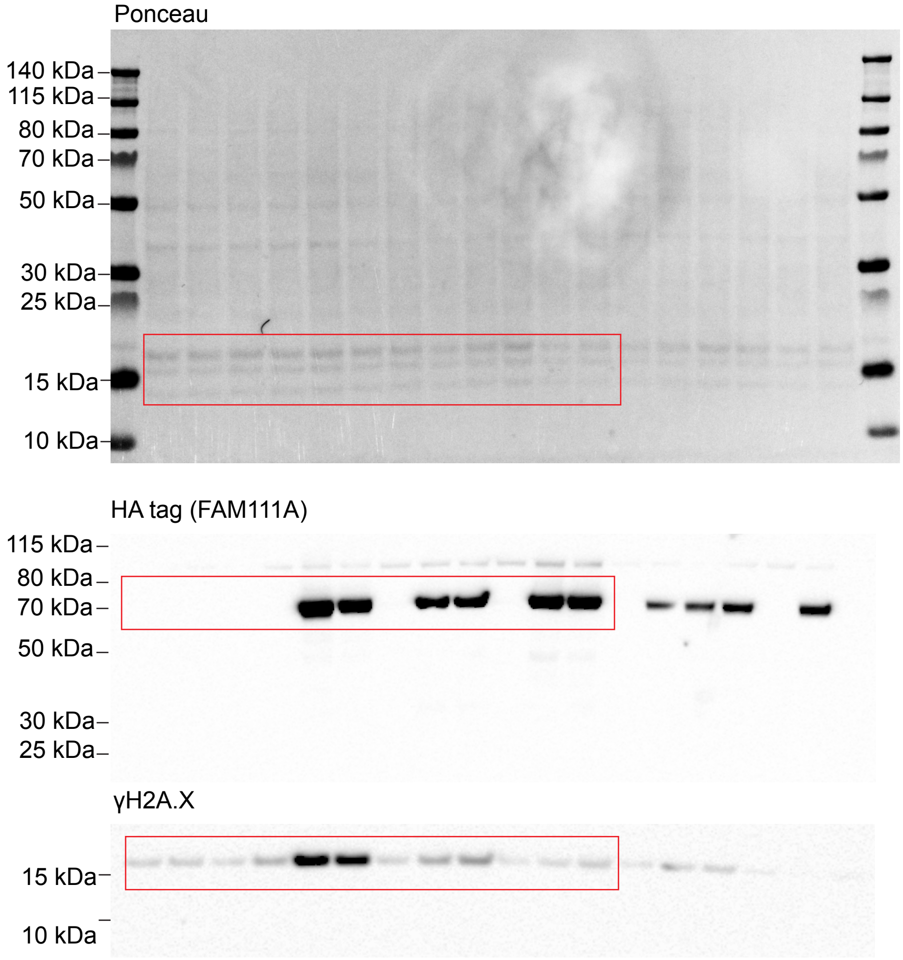


**Fig. S3G**

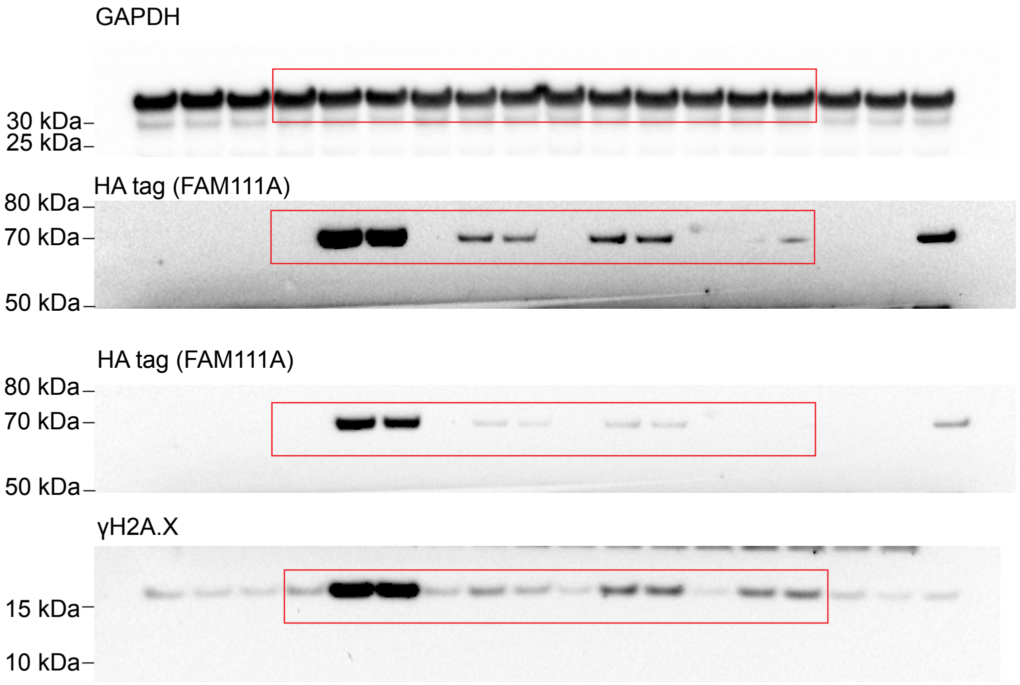

### Fig. S4B

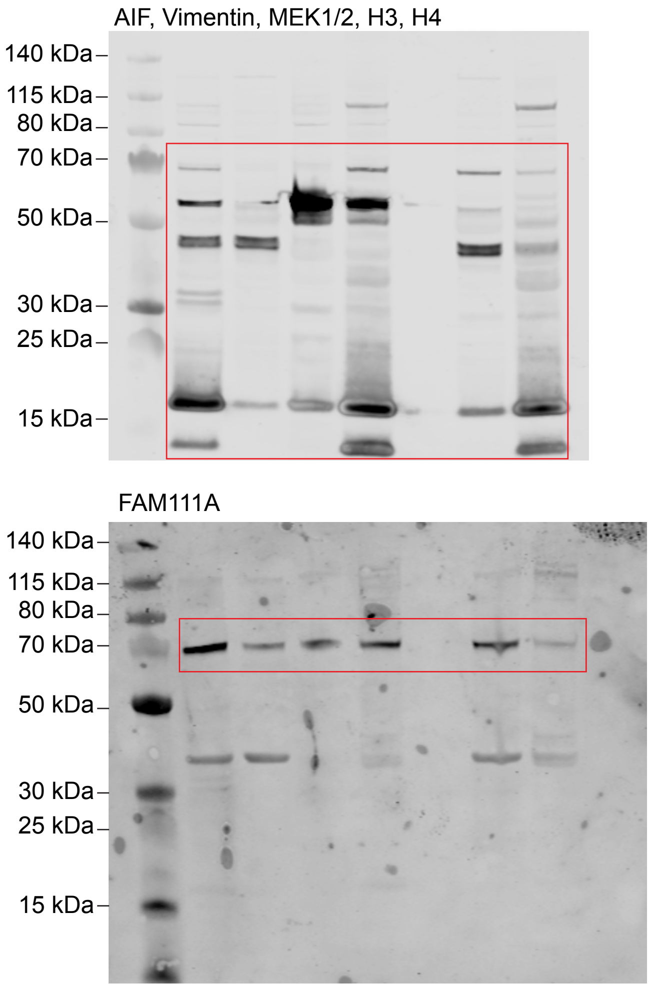


### Fig. S4C

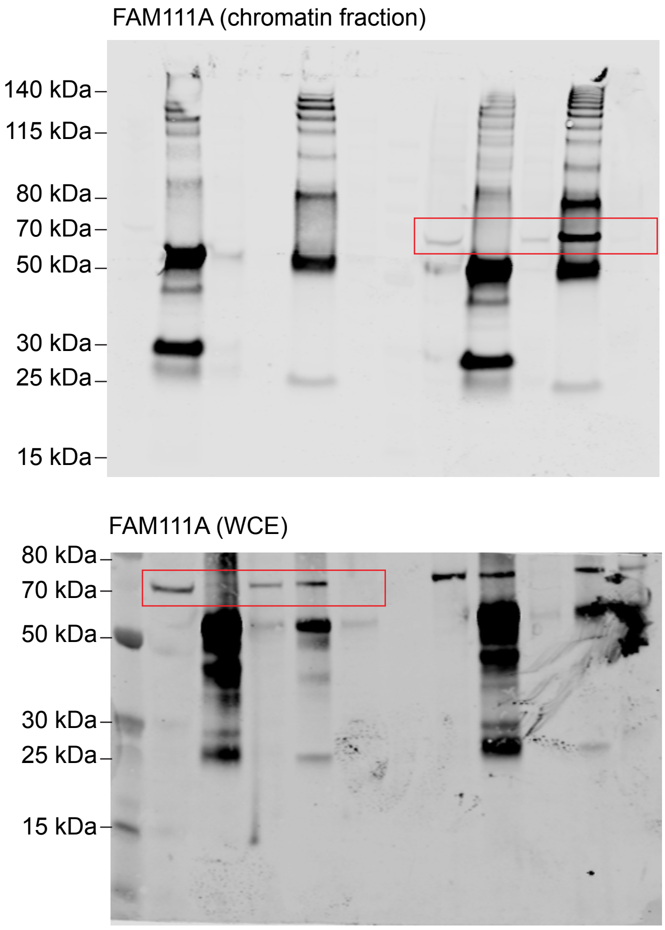


### Fig. S4D

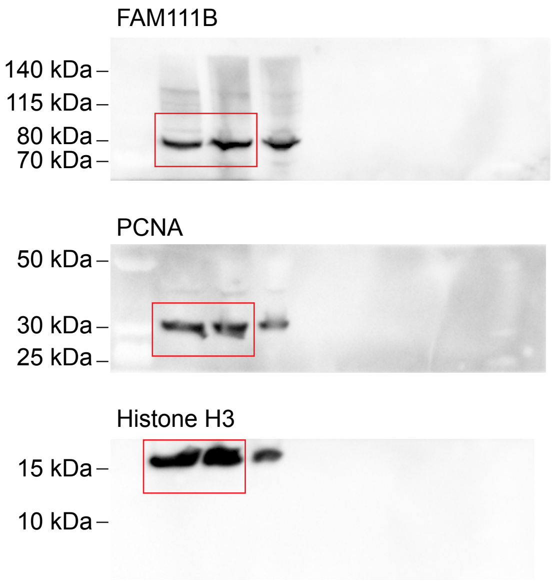


### Fig. S4F

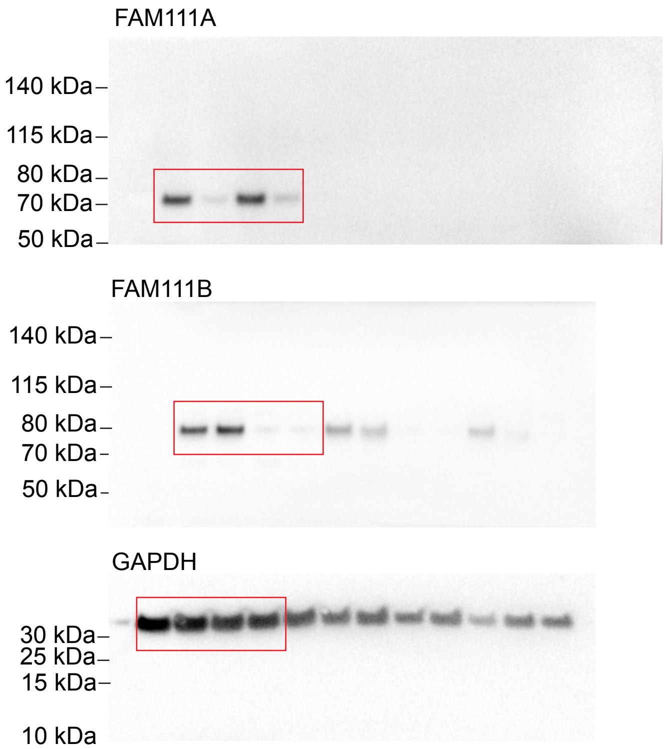
